# In-situ Single-Molecule Investigations of the Impacts of Biochemical Perturbations on Conformational Intermediates of Monomeric α-Synuclein

**DOI:** 10.1101/2023.08.31.555667

**Authors:** Wenmao Huang, Shimin Le, Mingxi Yao, Yi Shi, Jie Yan

**Affiliations:** Department of Physics, National University of Singapore, Singapore 117542; Mechanobiology Institute, National University of Singapore, Singapore 117411; Centre for Bioimaging Sciences, National University of Singapore, Singapore 117546; Research Institute for Biomimetics and Soft Matter, Fujian Provincial Key Lab for Soft Functional Materials Research, Department of Physics, Xiamen University, Xiamen, China 361005; Department of Biomedical Engineering, Southern University of Science and Technology, Shenzhen, China 518055; School of Chemistry and Molecular Engineering, East China Normal University, Shanghai, China 200241

## Abstract

α-Synuclein aggregation is a common trait in synucleinopathies, including Parkinson’s disease. Being an unstructured protein, α-synuclein exists in several distinct conformational intermediates, contributing to both its function and pathogenesis. However, the regulation of these monomer conformations by biochemical factors and potential drugs has remained elusive. In this study, we devised an in-situ single-molecule manipulation approach to pinpoint kinetically stable conformational intermediates of monomeric α-synuclein and explore the effects of various biochemical factors and drugs. We uncovered a partially folded conformation located in the non-amyloid-β component (NAC) region of monomeric α-synuclein, which is regulated by preNAC region. This conformational intermediate is sensitive to biochemical perturbations and small-molecule drugs that influencing α-synuclein’s aggregation tendency. Our findings reveal that this partially folded intermediate may play a role in α-synuclein aggregation, offering fresh perspectives for potential treatments aimed at the initial stage of higher-order α-synuclein aggregation. The single-molecule approach developed here can be broadly applied to the study of disease-related intrinsically disordered proteins.

## Introduction

Parkinson’s disease (PD) is a prevalent neurodegenerative disorder that affects approximately 10 million people around the world^1,2^. The cytopathological hallmarks of PD and other synucleinopathies include Lewy bodies and Lewy neurites, which contain aggregates of α-synuclein, a protein abundantly expressed in the central nervous system^3–5^. Several mutations in the α-synuclein encoding gene (SNCA) have been identified as key factors in the pathogenesis of PD, which modulate the aggregation propensity of α-synuclein^6–9^. Emerging evidence suggests that the misfolded α-synuclein monomers, aggregated cytotoxic oligomers, and small amyloid fibrils that act as prion-like seeds for aggregate propagation are the key pathological factors in synucleinopathies^5,10–16^.

α-synuclein is an intrinsic disordered protein (IDP) that has long been considered as a soluble monomer^17–19^. This 140-amino acids protein comprises three distinct regions (Fig. 1a): a positively charged N-terminal region (amino acid, a. a. 1−60), a non-amyloid-β component (NAC) region (a. a. 61−100,), and a negatively-charged intrinsic disordered C-terminal region (a. a. 101−140)^17,18^. We note that in our work, we expand the previously defined NAC region^18^ from 61−95 to 61−100, which constitutes the major core structure of α-synuclein amyloid fibril^20–24^. The hydrophobic core of α-synuclein aggregates is formed by the NAC region^25,26^, whose aggregation propensity is regulated by an N-terminus subregion (preNAC, a. a. 36−60, containing most of the mutations) adjacent to the NAC region^26,27^. The charged N-terminal and C-terminal regions exhibit transient long-range intra-molecule interactions as well as interactions with the cytoplasm components^17^, which are believed to help shield the hydrophobic NAC core from exposure to the cytoplasm and keep the monomer isoform in vivo^17,28,29^. Upon binding to the lipid membrane, the N-terminal and NAC regions can form an α-helical structure^30^, which represents the only solved monomeric structure of α-synuclein to date^18^.

**Figure 1.**
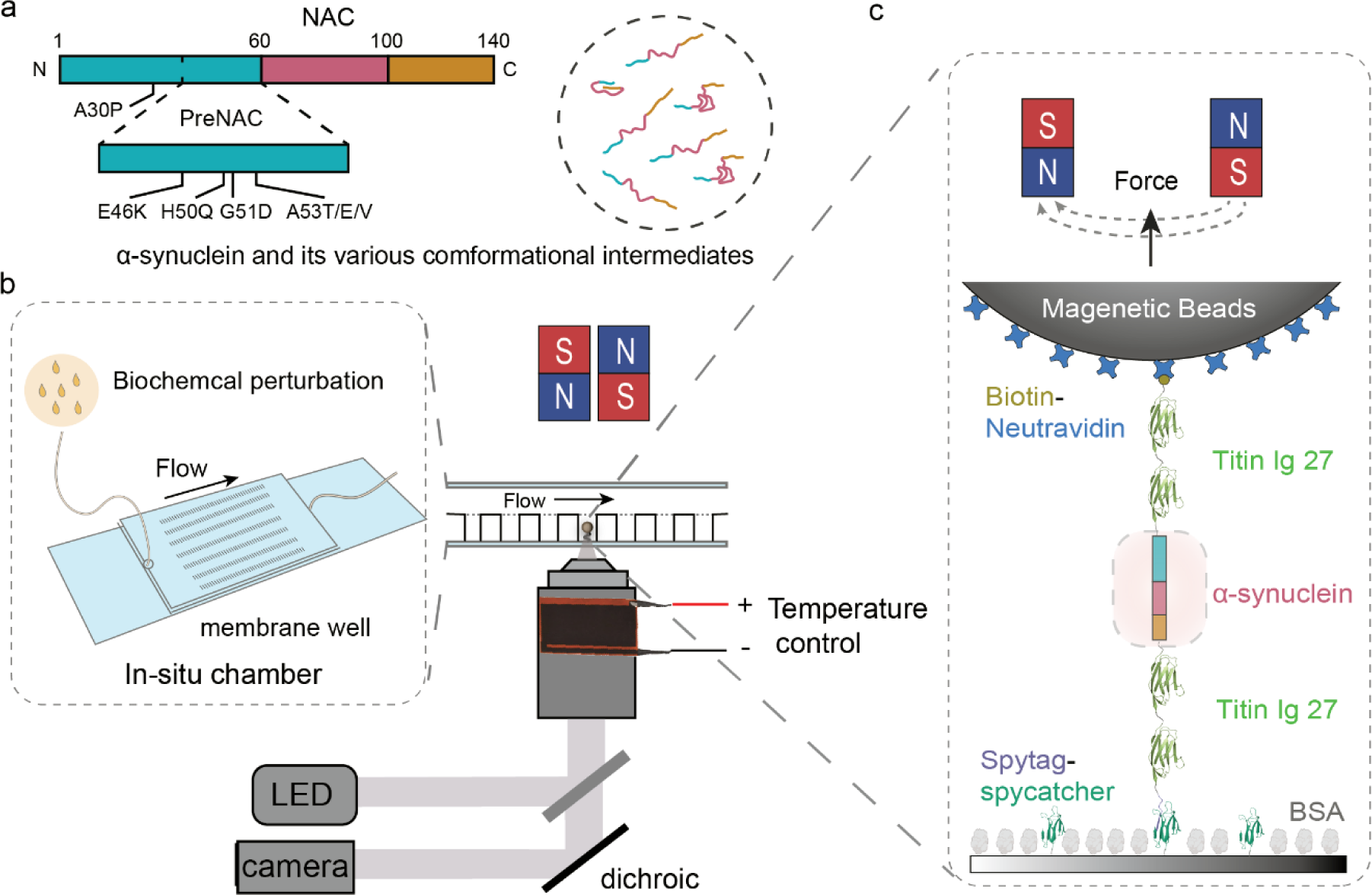
In-situ magnetic-tweezer-based single-molecule approach. (a) Illustrated sequence map of wild-type α-synuclein (1−140 a.a.), alongside potential conformational intermediates of α-synuclein. (b) Schematic diagram of the experimental setup for in-situ magnetic-tweezer-based single-molecule manipulation, comprising a magnetic tweezer for precise mechanical manipulation of proteins in the piconewton range with nanometer resolution, an in-situ buffer exchange chamber with membrane wells, and a temperature control system. (c) Elaborate schematics showcasing that the recombinant monomeric α-synuclein was tethered to the glass substrate and the superparamagnetic beads through specific SpyTag-SpyCatcher chemistry and biotin-neutravidin bond, respectively.

Cryo-electron microscopy has fully characterized the fibril-like structures of α-synuclein, revealing common fibril polymorphs featuring a β-rich kernel structure comprising the preNAC and NAC regions (36−100)^20–22,24,31^. During the propagation process from soluble monomers to fibril-like aggregates, cytotoxic^11,12,14,32^ or protective^11,33^ oligomers have been proposed to exist. At the monomer level, increasing evidence suggests that the disordered monomer is highly dynamic with versatile conformational structures (Fig. 1a), which can be influenced by various physiologically and pathologically relevant conditions^17,18,28,29,34–36^. In-depth in-vitro or in-vivo nuclear magnetic resonance (NMR) experiments have systematically studied the inter- or intramolecular interactions of α-synuclein, including the effects of lipid vesicle binding, N-terminal and C-terminal region regulation, chaperone binding, post-translational modifications, and mutations^17,18,27–29,35,37^.

Importantly, circular dichroism (CD) and NMR spectral studies have suggested the presence of partially folded intermediates of α-synuclein with varying aggregation propensity^17,18,28,34^. Some of the intramolecular interactions have reported to suppress the aggregation of α-synuclein, where the exposed NAC is shielded by long-range intramolecular interactions involving the NAC and C terminus^38,39^. Consistently, several single-molecule studies have reported multiple conformational intermediates in monomeric α-synuclein^36,40–43^. In particular, an optical tweezers-based study reported that the majority (85%) of the sampled α-synuclein monomers lacked any stable or metastable structures, while the remaining 15% contained diverse partially folded structures that can unfold at forces within 15 pN^36^.

Building on previous reports of intermediates in monomeric α-synuclein, along with their potential connection to aggregation, we embarked on an investigation into the regulation of conformational intermediates in monomeric α-synuclein. We utilized a magnetic-tweezer-based single-molecule manipulation approach for this purpose, where the potential intermediates in monomeric α-synuclein were recognized based on their unique mechanical signatures. We then applied in-situ perturbations to examine the effects on these intermediates. In line with previous research, we identified multiple conformational intermediates. Besides a primary conformational intermediate, characterized by a disordered and randomly coiled peptide conformation, we uncovered a previously unknown major intermediate that features a partially folded element in the NAC region with a stable structure. This intermediate, henceforth referred to as the PF intermediate, is regulated by the preNAC region. Intriguingly, and potentially significantly, the PF intermediate can be finely adjusted through biochemical perturbations and specific potential drugs known to influence higher-order α-synuclein aggregation.

## Results

### In-situ single-molecule manipulation approach

To detect potential conformational intermediates in monomeric α-synuclein, we developed a magnetic-tweezer-based single-molecule approach to probe its mechanical signatures. Specifically, α-synuclein was spanned in four well-characterized titin I27 Ig domains^44^, which served as a molecular spacer and mechanical fingerprint for single-molecule tethers (Supplementary Note 1). It has been reported that spanning α-synuclein between structural domains can effectively slow down higher-order oligomerization of α-synuclein^45^. Consistently, we confirmed that the spanning I27 domains in our study prevented aggregation or oligomerization of the recombinant protein constructs (Supplementary Fig. 1). Moreover, we found that the recombinant proteins remained as monomers for up to five days (Supplementary Fig. 1, Supplementary Note 1).

The mechanical signatures of potential distinct conformational intermediates of α-synuclein could be manifested as incremental extension changes resulting from abrupt conformational shifts, along with the corresponding forces when these transitions occur. To examine how biochemical factors and temperature impact the conceivable conformational intermediates of monomeric α-synuclein, we employed a disturbance-free membrane-well system that mitigates unwanted flow-induced stretching of the single-molecule tether during buffer exchange^46^ (Fig. 1b). This system is crucial for the in-situ analysis of the effects of various biochemical perturbation. Furthermore, to investigate the potential influence of temperature on conformational intermediates, we employed an objective-heating system to precisely regulate the temperature within the sample chamber (Fig. 1b).

The recombinant protein construct, which was fused with an N-terminal biotinylated AviTag and a C-terminal SpyTag, was anchored between a Neutravidin-coated superparamagnetic microbead (2.8 µm in diameter, M270, Invitrogen) and a SpyCatcher-coated coverslip glass surface (see Fig. 1c). Subsequently, this tether was subjected to cycles of force-increase and decrease scans between 1.5 ± 0.1 pN and 100 ± 10 pN at constant loading rates (see Supplementary methods). We would like to note that the transition forces are associated with a 10% uncertainty arising from our force-calibration method^47^. During these force scans, the height of the end-attached superparamagnetic microbead was continuously monitored. We note that when there is a change in force, the corresponding change in bead height is influenced by two factors: the change in the extension of the tethered molecule and the rotation of the bead due to torque rebalancing^47^. Conversely, bead height changes at a constant force or stepwise bead height changes during force loading represent the extension change of the molecule. The specific mechanical responses of a single tethered α-synuclein molecule were ensured by the signatures of mechanical unfolding of the four I27 domains at forces above 80 pN (Supplementary Fig. 2).

### Identification of conformational intermediates of monomeric α-synuclein

We demonstrate a significant conformational species (disordered) of monomeric α-synuclein, as evidenced by a smooth force-bead height curve that lacks stepwise structural transitions over the experimental time scale at specific loading rates (Fig. 2a, Supplementary Fig. 2a). Due to the aforementioned bead rotation, the smooth force-bead height curve does not represent the force-extension curve of the tethered molecule. Consequently, it is not possible to estimate the contour length of the underlying disordered polypeptide chain. However, our supplementary control experiments strongly suggest that the observed smooth force-bead height curves correspond to a completely random, unstructured coiled conformation of α-synuclein (Supplementary Note 2 and Supplementary Fig. 3). This aligns with the anticipated presence of such an unstructured coiled conformation given the inherently disordered nature of this protein, and is consistent with the observation from previous studies.

**Figure 2.**
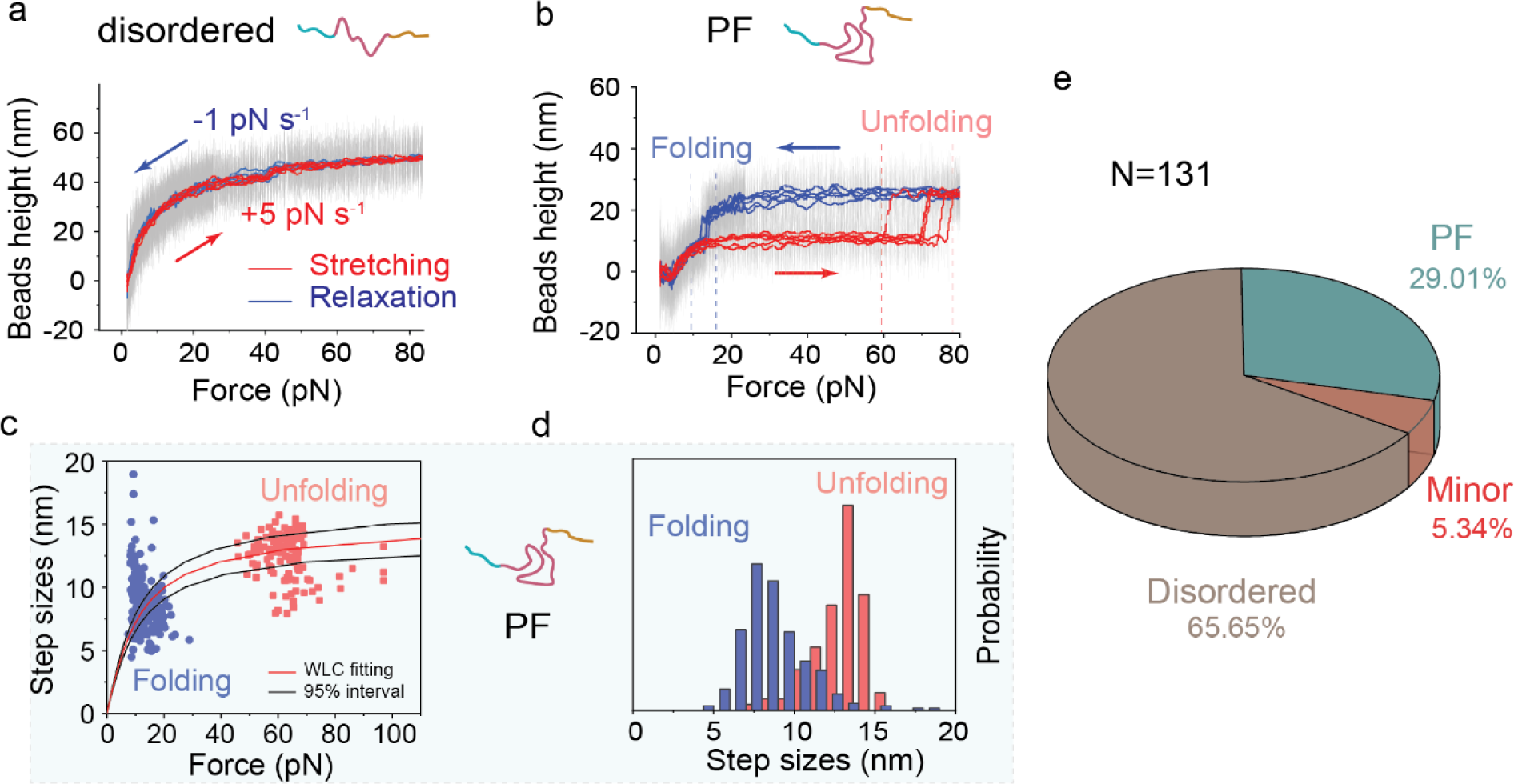
Two major conformational intermediates of monomeric α-synuclein. (a) Typical force−bead height curves of a disordered monomeric α-synuclein during force-increase scan (red, loading rate of 5 pN s^−1^) and subsequent force-decrease scan (blue, loading rate of −1 pN s^−1^). (b) Typical force−bead height curves of partially folded (PF) α-synuclein during force-increase and force-decrease scans at the same corresponding loading rates as in (a). (c) Unfolding and refolding force−step size distribution of PF α-synuclein at loading rates of 5 pN s^−1^ (red dots) and −1 pN s^−1^ (blue dots), respectively. A Worm-like chain model was used to fit the data (red line), and the optimal fitting persistence length (A) and contour length (Lc) were obtained as A= 0.50 ± 0.05 nm, Lc= 15.50 ± 1.50 nm. ± indicates S.E.. (d) Normalized unfolding and refolding step size distributions of PF α-synuclein, respectively. The data in (c) and (d) are from 124 (unfolding) and 268 (refolding) events, which were collected from more than 30 independent α-synuclein tethers. (e) Conformation fractions of monomeric α-synuclein (N=131 molecules) based on their distinct mechanical signatures (Supplementary Fig. 8).

Unexpectedly and intriguingly, the experiments also revealed the existence of another major conformational intermediate, i.e., the PF intermediate, featured with a partially folded element. During force-increase scans at a loading rate of 5 pN s^−1^, this conformational species exhibited a distinct unfolding step at approximately 60 pN associated with a step size of around 14 nm. Subsequently, when force was reduced at a loading rate of −1 pN s^−1^ to around 15 pN, this species could refold, resulting in a step size of roughly 8 nm (Fig. 2b−d and Supplementary Fig. 2b). The unfolding and refolding step sizes were indicative of the release or absorption of a peptide of 41 ± 4 amino acids (a.a.) under force (Fig. 2d and Supplementary Fig. 4). This uncertainty contributes to the spread of the step size histograms, similar to the level observed in our previous studies of well-defined structural domains, such as titin I27^44^ and α-actinin spectrin repeats^48^. This peptide (41 ± 4 a.a.) is much shorter than α-synuclein’s total amino acid count (140 a.a.), implying that the unfolding and refolding signatures are from partially folded element in the PF intermediate. Additionally, the PF intermediate was detected in control experiments using the α-synuclein protein without the I27 domains, which rules out the possibility that it is induced by the spanning I27 domains (Supplementary Fig. 5 and Supplementary Note 1).

The partially folded element in the PF intermediate exhibits a high level of mechanical stability. It required high forces (typically > 50 pN) for unfolding at various force-increase loading rates (0.5 − 5 pN s^−1^) (Supplementary Fig. 4). It also survived substantial lifetime at constant high forces (40 − 60 pN, Supplementary Fig. 6, Supplementary Note 3 for force-jump experiments). Over 60% of the unfolded intermediate rapidly refolds at forces of around 15 pN at various force-decrease loading rates (−2.0 − −0.1 pN s^−1^) (Supplementary Fig. 4). The rapid refolding of the partially folded intermediate is surprising and noteworthy, as it was not observed in disordered conformations. This implies that the unfolded intermediate’s conformation differs from that of entirely disordered species and may lead to another intermediate that can refold back when forces are reduced to 15 pN. Although the unfolded PF species’ conformation remains unclear, the outcome suggests that it may be a force-resistant structural intermediate that requires further investigation. Additionally, we found no conversion between the disordered and PF intermediates at 23 °C within our experimental time scale (tens of hours), indicating that they represent local energy minima separated by a high free energy barrier. Based on these observations, we note that the stable partially folded intermediate in the PF species is different from the intermediates reported in previous single-molecule manipulation studies^36,40–43,49^.

In addition to the primary conformational species (disordered: approximately 65%; PF: approximately 30%), we also identified a minor conformational species (roughly 5%) consisting of various partially folded intermediates. Each of these intermediates exhibits distinct mechanical signatures based on transition forces and step sizes (refer to Supplementary Note 4 and Supplementary Fig. 7). The population fractions of these conformational species (Fig. 2e and Supplementary Fig. 8) have been quantified using the mechanical signatures from randomly selected tethered protein constructs, which represent the population fractions of different α-synuclein conformational species coexisting in the solution.

### The partially folded element situates in NAC and is regulated by preNAC

To locate the region of the partially folded intermediate in the PF species and the regions that may influence its stability, we carried out single-molecule experiments on various truncated α-synuclein constructs (Fig. 3a), including NAC (a.a. 61−100), NAC + preNAC (36−100), ΔC (1−100), and three N-terminal truncations ΔN1 (11−140), ΔN2 (36−140), ΔN3 (61−140), as well as two truncates termed as ΔNAC, ΔpreNAC without NAC or preNAC region. Like wild-type α-synuclein, all truncated α-synuclein variants were spanned between two I27 domains at each side to maintain their monomeric conformations (See Supplementary Information for details of protein construct sequences).

**Figure 3.**
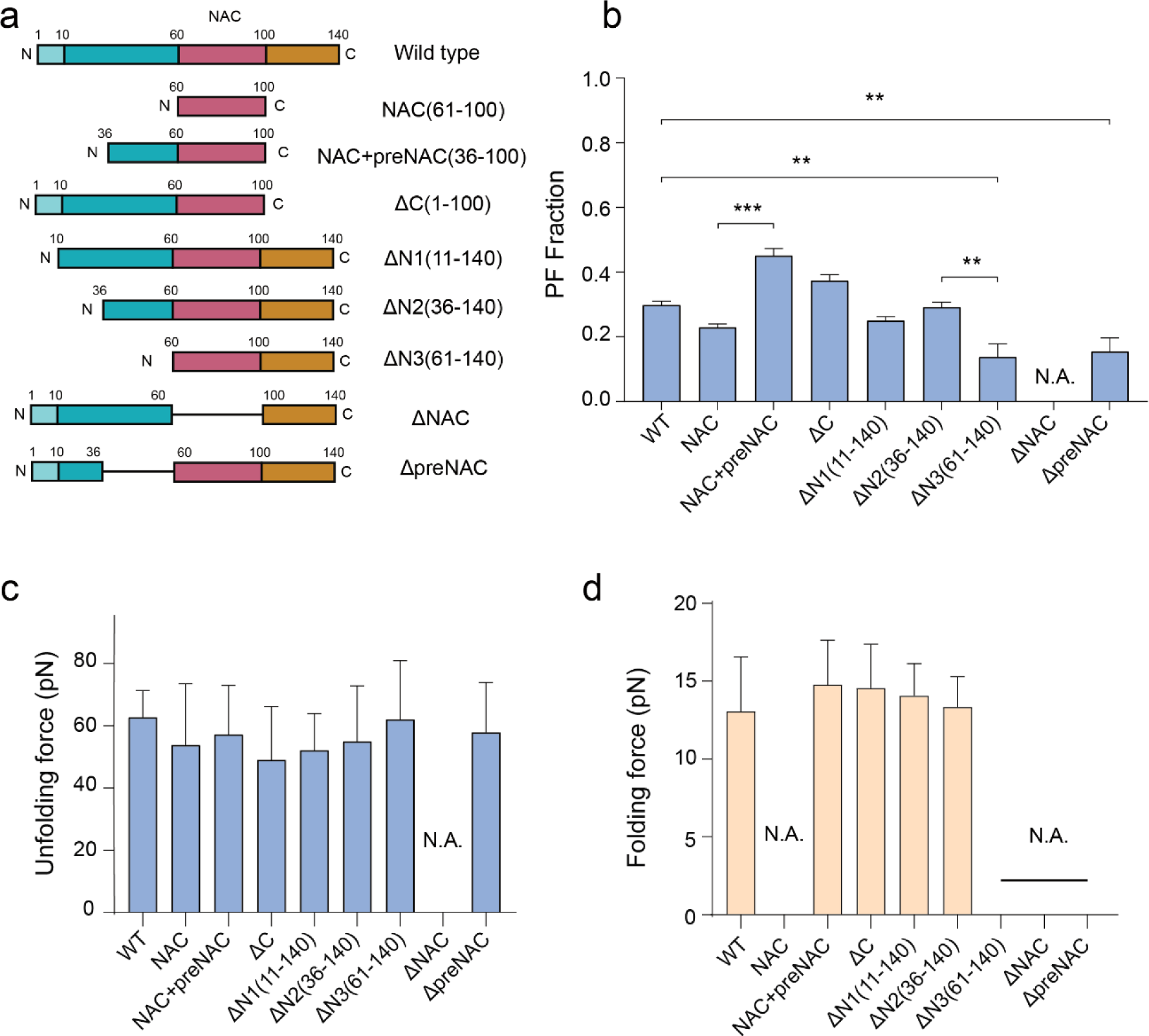
Effects of truncations of α-synuclein. (a) Illustrated domain maps of wild-type and truncated α-synucleins. (b) PF population fractions of wild-type α-synuclein (WT) and various truncations based on the unfolding signature of the PF species. The numbers of tested molecules are 131, 35, 20, 74, 60, 48, 51, 47, and 52 for WT, NAC, NAC+preNAC, ΔC, ΔN1, ΔN2, ΔN3, ΔNAC, and ΔpreNAC, respectively. Error bars indicate mean ± S.E.M.. (c) The average unfolding forces during force-increase scans at a loading rate of 5 pN s^−1^. The numbers of unfolding events are: 124, 153, 193, 199, 161, 237, 16, 0, and 21 for WT, NAC, NAC+preNAC, ΔC, ΔN1, ΔN2, ΔN3, ΔNAC, and ΔpreNAC, respectively. (d) The average folding forces during force-decrease scans at a loading rate of −1 pN s^−1^. The numbers of folding events are: 268, 0, 76, 55, 98, 89, 0, 0, and 0 for WT, NAC, NAC+preNAC, ΔC, ΔN1, ΔN2, ΔN3, ΔNAC, and ΔpreNAC, respectively. Error bars in Fig. 3c, 3d indicate mean ± S.D.. N.A. means not applicable (no PF conformation is observed). **: p < 0.01; ***: p < 0.001 (95% of confidence intervals). p values were determined using an unpaired two-tailed Student’s t-test.

The primary findings from single-molecule force scans are summarized in Fig. 3b and Supplementary Fig. 9. The complete disappearance of the characteristic unfolding/refolding signatures of PF was observed in the NAC truncation, indicating that the partially folded intermediate is in the NAC region. Apart from the ΔNAC construct, all other truncated variants retained the characteristic unfolding signature of the partially folded intermediate.

The unfolding and refolding signatures of the PF species for NAC+preNAC, ΔC, ΔN1, and Δ N2, were found to be identical to those of wild-type α-synuclein (Figs. 3c−d, Supplementary Figs. 10−11). Notably, in the absence of the preNAC region in NAC, ΔN3, and ΔpreNAC, the unfolding signature was preserved, whereas the refolding only occurred at ∼ 1.5 pN at a loading rate of −1 pN s^−1^ (Fig. 3d, Supplementary Figs. 10−11). The refolding of the unfolded structure undergoes either the rapid refolding pathway characterized by the ∼15 pN refolding forces or the slow refolding pathway at ∼ 1.5 pN (Supplementary Fig. 11c). The presence of the preNAC region was found to be essential to the rapid refolding pathway, while its absence led to a much slower refolding of the unfolded PF structure (Supplementary Figs. 11d−f). These results suggest that the preNAC region modulates the refolding kinetics of the PF’s partially folded intermediate after its unfolding.

Based on these observations, we conclude that 1) the NAC region bears the identified partially folded element in the PF intermediate, and 2) the preNAC region facilitates its folding thus contributing to the regulation of the partially folded intermediate. We speculate that preNAC might interact with the unfolded conformation of the element, promoting folding along a specific pathway. Notably, prior studies have reported that the deletion of preNAC significantly suppresses the aggregation of α-synuclein^27^, while several familial mutations of early onset PD were located in the preNAC region^6–9^. We reason that these mutations may influence the preNAC-mediated regulation of the PF species, which is our ongoing study.

### Biochemical perturbations on PF intermediates

Given that the identified PF intermediate was not reported in previous studies and that it represents a major form of the monomeric α-synuclein, we investigated the potential effects of two known aggregation promoting factors, temperature and pion-like seeds, on the formation of the PF intermediate.

It is known that the preparation of α-synuclein can have a pronounced effect on its aggregation properties^50^. In order to ensure identical initial conditions for the investigations, the monomeric α-synuclein proteins were fully unfolded using 6 molars (M) guanidine hydrochloride (GuHCl) and 8 M urea, and subsequently diluted and flowed into the in-situ sample chamber to form single-molecule tethers. Treatment with 6 M GuHCl and 8 M urea resulted in a dramatically decreased fraction of the PF intermediate to a negligible level within hours after the diluted denaturant solution was removed from the chamber, leaving the disordered conformation as the predominant species (Supplementary Note 5 and Supplementary Fig. 12). This finding is consistent with the ability of the selected denaturants (GuHCl and urea) to fully convert monomeric α-synuclein into an unstructured conformation, regardless of its initial state following protein expression and purification.

To investigate the effect of temperature, a tethered α-synuclein protein in the disordered conformation was identified at 23 °C and in-situ subject to a temperature increase to 37 °C within 10 minutes (Fig. 4a). The conversion to the PF intermediate of the same α-synuclein molecule was observed within hours at 37 °C (Fig. 4b), while such conversion was not observed within 10 hours at 23 °C. This result indicates that higher temperatures increase the rate of the formation of the PF intermediate converted from an initially disordered conformation. In the experiment to test the effect of prion-like seed perturbation, a tethered α-synuclein protein in the disordered conformation was identified at 23 °C and then in-situ exposed to a solution containing prion-like seeds (∼10 nM, equivalent to the molar concentration of monomer) at the same temperature (Fig. 4a). The conversion of the same α -synuclein molecule to the PF intermediate conformation was also observed within hours after the introduction of seeds (Fig. 4c). In contrast, without the seeds, such conversion was not observed up to tens of hours at 23 °C (Fig. 4d).

**Figure 4.**
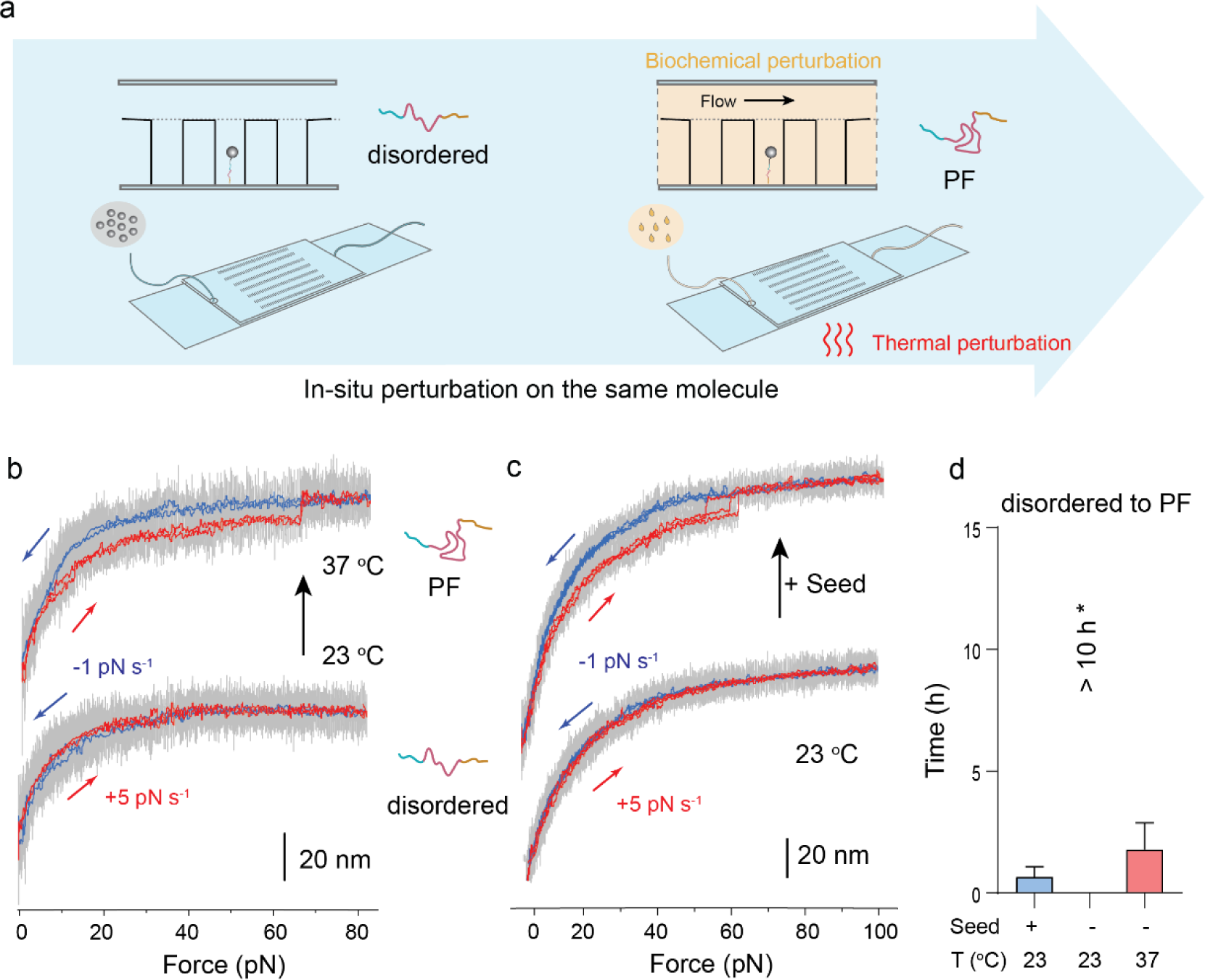
In-situ perturbations of the disordered-to-PF conversion. (a) Schematic illustration of the in-situ single-molecule experiments. Single disordered α-synuclein was identified at 23 °C (left panel), and the seed (small α-synuclein aggregates) was introduced or the temperature was increased to 37 °C (right panel). (b) Representative force−bead height traces of the disordered α-synuclein at increasing temperatures from 23 °C (bottom) to 37 °C (top). Similar results were observed in 3 independent tethered proteins. (c) Representative force−bead height traces of the α-synuclein before (bottom) and after (top) flowing in 10 nM α-synuclein seed at 23 °C. Similar results were observed in 4 independent tethered proteins. (d) Average times required for the conversion from the disordered to PF conformations of α-synuclein under the corresponding in-situ perturbations. Error bars indicate mean ± S.E.M.. * For comparison, the disordered-to-PF transition at 23 °C without seed was not observed over 10 hours.

In the above-mentioned in-situ experiments, the conversion from the disordered species to the PF intermediate, was based on similar unfolding forces and step sizes to those observed in Fig. 2. However, we observed a difference that the previously noted refolding of the unfolded PF at forces around 15 pN is absent in both cases (higher temperature or seed treatment). This could indicate an effect from temperature and seed on its refolding pathway. In addition, we found that the disordered α-synuclein could also be converted to PF intermediate by in-vitro pre-treatment experiment, where the denatured unstructured proteins in solution were treated by seeds or incubated at different temperature, prior to single-molecule testing (Supplementary Note 5, Supplementary Fig. 12).

Together, the in-situ perturbation single-molecule mechanical assays have shown that the PF intermediate of the monomeric α-synuclein molecules is positively modulated by increased temperature and prion-like seeds.

### Two potent small-molecule inhibitors of PF

Several natural or synthesized small molecules have been demonstrated to suppress the formation of α-synuclein fibrils, and some of these compounds are potential therapeutic agents for neurodegeneration diseases associated with α-synuclein aggregation^51–57^. Multiple mechanisms have been proposed to explain the activity of these small molecules, including destabilization of the monomer−monomer interface within α -synulcein fibrils and modification of the conformation of α-synuclein monomers to discourage aggregation^51,53,54,57,58^. Here, we explore the possibility that certain small molecules may affect the PF intermediate of monomeric α-synulcein identified in this study.

The recombinant α-synuclein proteins were pre-treated with six small molecules (Fig. 5a) at a concentration of 0.1 mM for 24 hours at 23°C, without prior exposure to a denaturant. After removing the small molecules from the single-molecule experimental flow chamber, we assessed the initial population fraction of the PF intermediates using magnetic tweezers. Four of the six small molecules, namely dopamine, baicalein, curcumin, and fasudil, showed significant suppressive effects on the PF species. Dopamine and baicalein exhibited the most potent inhibitory effects (Fig. 5b). As α -synulcein monomers were not pre-treated with a denaturant, it suggests that the potent inhibitory effects of dopamine and baicalein are mediated by direct binding and catalyzing conversions from the PF species to other conformations.

**Figure 5.**
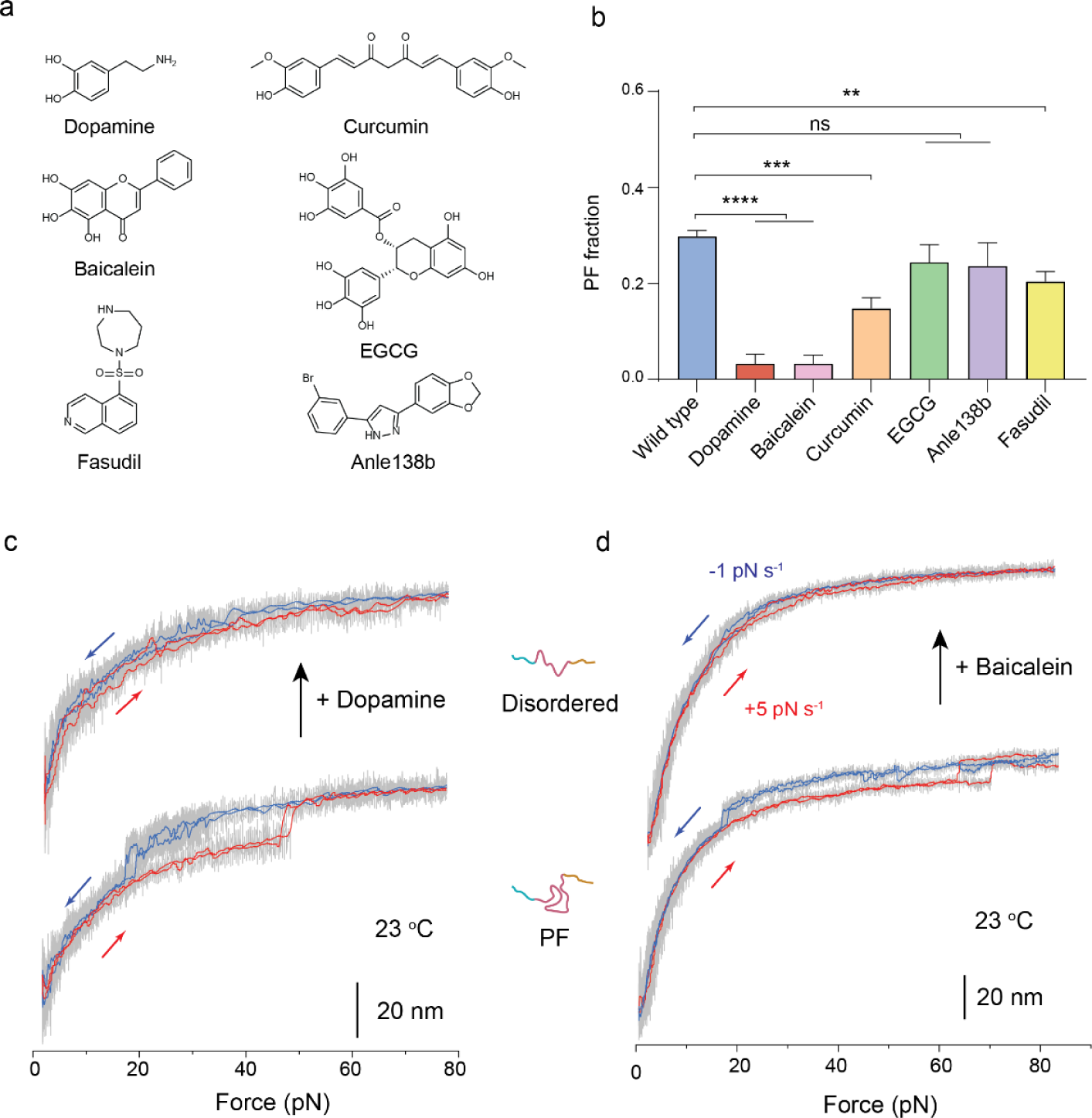
Effects of small-molecule drugs. (a) Chemical structures of six small-molecule drugs reported to inhibit the in-vitro aggregation of α-synulcein. (b) PF fraction of wild-type and small-molecule pre-treated α-synulcein. The total molecule numbers are N=131, 62, 61, 61, 45, 55, and 54 for wild type, six small-molecule (dopamine, baicalein, curcumin, EGCG, anle138b, and fasudil) treated α-synulcein, respectively. Error bars indicate mean ± S.E.M.. ns: p > 0.05; **: p < 0.01; ***: p < 0.001; ****: p < 0.0001 (95% of confidence intervals). p values were determined by unpaired two-tailed Student’s t-test. (c−d) In-situ analysis of the PF-to-disordered transition by dopamine and baicalein. (c) Representative force−bead height curves of a PF α-synuclein before (bottom) and after (top) introducing 0.1 mM dopamine solution. Similar results were repeated on 3 independent α-synucleins. (d) Representative force-bead height curves of a PF α-synuclein before (bottom) and after (top) introducing 0.1 mM baicalein solution. Similar results were observed on 5 independent tethered proteins.

The effects of dopamine and baicalein were further confirmed with in-situ single-molecule experiments on the same tethers. A tethered α-synuclein protein in the PF intermediate conformation was identified at 23 °C and then in-situ exposed to a solution containing 0.1 mM dopamine or baicalein at the same temperature. The conversion of the same α -synuclein molecule to from the PF to disordered conformation was observed within 30 minutes after the introduction of the drugs (Figs. 5c−d).

Previous studies have demonstrated that dopamine can directly bind α-synuclein and might alter its conformation^53,59^, yet details of the inhibitor’s binding site and whether they could bind to the unfolded state^60^ remain unclear. Based on our observation that dopamine and baicalein exhibit potent inhibitory activity against the PF conformation, we reason that dopamine and baicalein may bind to critical residues in pre-NAC/NAC regions that suppresses the PF conformation. PD has been linked to low or declining levels of dopamine in the brain^59,61^. Our results provide a possible mechanistic understanding, as decreased dopamine level may lead to increased PF species that could promote α-synuclein aggregation.

## Discussions

In this study, we have developed a single-molecule manipulation approach to delve into the distinct conformational states within monomeric α-synuclein molecules. More importantly, we have explored the modulation of the in-situ conversion among these conformational intermediates under various biochemical conditions. This single-molecule approach is integrated with conventional bulk assays, providing a comprehensive comprehension of the diverse conformational landscape of monomeric α-synuclein.

The method we propose involves the generation of monomeric α-synuclein molecules by spanning them between two titin I27 immunoglobulin domains situated at each end. Biochemical assays have substantiated the ability of these I27 domains to sustain the monomeric state of α-synuclein for a duration of at least five days. Additionally, our methodology facilitates the discrimination of multiple conformational states of monomeric α-synuclein, discerned by unique mechanical signatures. Specifically, these signatures encompass unfolding and refolding forces, as well as transition step sizes observed during cycles of incremental and decremental force scans, quantified using a magnetic tweezers setup. The in-situ conversion of monomeric α-synuclein is facilitated through a temperature control system paired with a flow disturbance-free rapid solution exchange technology. Collectively, these techniques offer a robust, magnetic tweezers-based single-molecule platform ideal for investigating the in-situ conversions of various conformational states of monomeric α-synuclein, and potentially other intrinsically disordered proteins.

One significant discovery from this study is the unveiling of a previously undiscovered conformational state within monomeric α-synuclein, referred to as the PF intermediate. Distinguished by a stable, partially folded structure, the PF intermediate exhibits distinct mechanical traits in the NAC region–a domain acknowledged for composing the hydrophobic core of α-synuclein fibrils. The appearance of the PF intermediate is intricately modulated by the preNAC region, which facilitates its reformation following the unfolding of the partially folded element within the PF intermediate. Intriguingly, increased temperatures and the presence of prion-like seeds were found to accelerate the transformation from the initial disordered conformation to the PF intermediate, as unveiled through in-situ perturbations in single-molecule investigations. In contrast, dopamine and baicalein were observed to expedite the reverse conversion from the PF intermediate back to the original disordered conformation. These findings serve to enrich our comprehension of the intricate conformational landscape of monomeric α-synuclein (Supplementary Fig. 13). It is worth noting that previous single-molecule assays have missed the identification of the PF intermediate. This could be attributed to insufficient forces (<40 pN) that could not cause its unfolding in an optical-tweezers-based study^13d^, or non-optimally positioned fluorophores that were not sensitive enough to detect the extension difference between the disordered and the PF species in single-molecule FRET assays^40,42^.

The crucial initial stage leading to α-synuclein aggregation remains to be determined. Various studies have proposed that a key step in pathogenic aggregation involves oligomers^12,14,15,32,62–65^. Additionally, other research suggests that the monomeric conformational structures of α-synuclein may play a critical role in the early stages of aggregation^17,18,28,41,66^. The strong correlation between the occurrence of the identified PF intermediate and factors that are known to regulate the aggregation propensity of α-synuclein suggest a positive relation of the PF intermediate to the aggregation propensity of α-synuclein. Since α-synuclein aggregation requires hydrophobic residues in the NAC region^38,39^, in the PF intermediate the necessary hydrophobic residues required for aggregation should remain accessible. Once incorporated into the aggregated state, further conformational changes of α-synuclein proteins can be expected.

Finally, we would like to emphasize that this study is limited to in-vitro experiments solely based on mechanical signatures. As a result, it does not offer insights into the potential structure of this intermediate species. Furthermore, while the findings reveal a positive correlation between the intermediate in the PF species and α-synuclein aggregation, its actual roles in α-synuclein aggregation remain to be determined in future studies.

## Supporting information

Supporting informations

## Methods

### Reagents and Materials

(3-aminopropyl)triethoxysilane (APTES), glutaraldehyde solution (70% in H2O), dithiothreitol (DTT), bovine serum albumin (BSA), urea, guanidine hydrochloride (GuHCl), Sodium dodecyl sulfate (SDS), (−)-Epigallocatechin gallate (EGCG), curcumin from Curcuma longa (Turmeric), fasudil hydrochloride, baicalein, Anle138b, dopamine hydrochloride, thioflavin T (ThT), L-ascorbic acid, ampicillin, isopropyl β-d-1-thiogalactopyranoside (IPTG), phenylmethylsulfonyl fluoride (PMSF), biotin, mineral oil, PBS, HEPES, and Tris buffer were purchased from Sigma-Aldrich. NeutrAvidin biotin-binding protein, Dynabeads™ M-270 Epoxy beads were purchased from Thermo Scientific. Recombinant human α-synuclein protein monomer and recombinant human α-synuclein protein aggregate (seeds) were prepared following the previous method^67^.

### Gene construction, protein expression, and purification

The human α-synuclein SNCA gene (residue 1-140, Uniprot P37840) was codon-optimized and synthesized for expression in *Escherichia Coli* as gBlocks gene fragments (Integrated DNA Technologies (IDT)) with suitable overhangs. The gene was fused with four tandem titin immunoglobulin domains (Ig27) of a pET151 Vector (ampicillin resistance) with 6×His-tag for purification, AviTag for biotinylation, and SpyTag using the NEB HiFi assembly strategy (New England Biolabs, MA, USA). The construction of SpyTag binding protein−SpyCatcher was referred to the previous works^48^. Truncated variants (NAC (a.a. 61−100), NAC+preNAC (a.a. 36−100), ΔC (a.a. 1−100), ΔN1 (a.a. 11−140), ΔN2 (a.a. 36−140), ΔNAC (a.a. 1−60+101−140), and ΔpreNAC (a.a. 1−36+61−140)) were introduced using the Site-Directed Mutagenesis kit (New England Biolabs, MA, USA). The complete sequences of all recombinant protein constructs used in this study are listed in the Supplementary Information (“Protein construction and sequence” section). All the α-synuclein-Ig27 fused protein constructs were expressed in *E. Coli* DE3 with biotin protein ligase (BirA) and purified using His-tag affinity column following the previous protocol^48,68^.

The full-length α-synuclein monomer (1−140) expression plasmid (pET21a-alpha-synuclein) was a gift from Michael J Fox Foundation MJFF (Addgene plasmid # 51486). The α-synuclein monomer was expressed and purified following the previous method^20,21^. The purified α-synuclein monomer was concentrated to 500 µM in fibril growth buffer (50 mM Tris, pH 7.0, 150 mM KCl, 0.02% NaN3) and incubated at 37 °C with agitation (900 rpm) in an incubator (Thermo Fisher). After 5 days, the fibrillar samples were diluted to a concentration of ∼100 µM (equal to the monomer concentration) and ultrasonicated for 10 min (1 s on, 1 s off) on ice to yield small size of α-synuclein aggregates (seeds). More details on protein expression and purification are listed in the Supplementary Methods.

### Magnetic-tweezer-based single-molecule essay

Single-molecule detection on α-synuclein proteins was achieved on a homemade magnetic tweezers (MT) setup as previously described^47,69^. To perform the experiments, the α-synuclein-I27 fused protein constructs were tethered between Neutravidin-coated magnetic beads and SpyCatcher-coated cover glass in membrane-well laminar flow in-situ chambers. The standardized solution used for the single-molecule experiments was composed of 20 mM HEPES, 50 mM KCl, 10 mM MgCl2, 1% (m/m) BSA, 1 mM DTT, pH = 7.4, unless otherwise specified. Details on the preparation of the chamber and microbeads, as well as the sample preparation, magnetic tweezer setup, constant loading rate scan, force jump experiment, and force calibration, have been previously published in our studies^48,68^ and are briefly summarized in the Supplementary Information.

### In-situ single-molecule experiments

In a typical in-situ single-molecule experiment, the chamber was connected to a buffer exchange system to facilitate changes in solution condition. The tethered protein was initially scanned for several cycles to obtain its structural conformation prior to the buffer exchange. Solutions containing denaturant, seeds, or drugs, totaling 200 µL (three times the chamber volume), were added to the rapid buffer exchange system and flowed through the chamber. The membrane-well based disturbance-free system would inhibit the unwanted flow stretching perturbation to the tethered molecule. After about 10 minutes of incubation, the system stabilized, and force scans of the α-synuclein in the new buffer condition began.

In a typical in-situ temperature-change single-molecule experiment, the temperature of the experiment was increased from 23 to 37 °C using an objective heating system (Bioptechs) that had been pre-calibrated with a thermometer. The tethered protein was initially scanned for several cycles at 23 °C to obtain its structural conformation prior to the temperature change. The temperature was then increased to 37 °C and maintained for more than 10 minutes to allow for temperature equilibrium.

### Single-molecule data analysis

Raw time traces data were recorded at a sampling rate of 200 Hz. In all the figures presented in the main text and supplementary information, the time traces and bead height−force traces were smoothed using the fast Fourier transform (FFT)-smooth function of OriginPro 9.0, unless otherwise mentioned. The time resolution of our experiment is around 5 microseconds due to the sampling rate. To determine the unfolding and refolding force and step size for α-synucleins at different force loading rates, we collected data from multiple independent tethers. The figures and legends indicate the number of unfolding/refolding events (n) and the number of tethers (N).

To measure the conformational species fraction of α-synuclein molecules, the tethered single-molecule α-synucleins were randomly selected and scanned for more than ten cycles to obtain the mechanical responses. As the different conformations were kinetically separated, the conversion between the disordered conformation and PF was not observed over the single-molecule experimental time scale (tens of hours) at 23 °C. Therefore, the tethered α-synuclein conformation fractions corresponded to the population fractions of the different conformational species of α-synuclein co-existing in the solution.

### Reporting summary

Further information on research design is available in the Nature Research Reporting Summary linked to this paper.

## Data availability

All data generated and analysed during this study are included in this article and supplementary information and are also available from the authors upon reasonable request.

## Acknowledgments

We thank the High-throughput Molecular Genetics core of the Mechanobiology Institute for the help with protein expression. This work was supported by the Singapore Ministry of Education Academic Research Fund Tier 2 (MOE-T2EP50220-0015), the Singapore Ministry of Education Academic Research Fund Tier 3 (MOET32021-0003), the Ministry of Education under the Research Centres of Excellence programme, and the National Natural Science Foundation of China (No. 21902075).

## Author contributions

J.Y. and W.H. conceived the project and designed the experiments. W.H., S.L. and M.Y. contributed to the methodology development. W.H., S.L. and Y.S. carried out the experiments and data analysis. J.Y. supervised the project. W.H. and J.Y. wrote the manuscript with contributions from all authors.

## Additional information

**Supplementary information** is available for this paper at ****.

**Reprints and permissions information** is available at www.nature.com/reprints.

**Correspondence and requests** for materials should be addressed to J.Y..

**Publisher’s note:** Springer Nature remains neutral with regard to jurisdictional claims in published maps and institutional affiliations.

