## Supporting informations for "In-situ Single-Molecule Investigations of the Impacts of Biochemical Perturbations on Conformational Intermediates of Monomeric α-Synuclein"

**Corresponding Author**

**This PDF file includes:**

Supplementary Methods

Supplementary Notes 1−5

Supplementary Figure 1. Bulk Characterizations

Supplementary Figure 2. The mechanical responses of wild-type monomeric α-synuclein-I27 fused protein.

Supplementary Figure 3. The mechanical responses of disordered α-synuclein.

Supplementary Figure 4. The mechanical responses of PF α-synuclein under loading rate force scan.

Supplementary Figure 5. Control experiments using the α-synuclein protein without the I27 domains.

Supplementary Figure 6. The mechanical responses of PF α-synuclein in force jumping experiments.

Supplementary Figure 7. The mechanical responses of the minor species of α-synuclein.

Supplementary Figure 8. 2D map summarizes the mechanical responses of monomeric α-synuclein.

Supplementary Figure 9. Species fractions of wild-type α-synuclein and its truncations based on the unfolding signatures.

Supplementary Figure 10. Typical force−bead height traces of α-synuclein truncations with PF mechanical signature.

Supplementary Figure 11. Mechanical signatures of truncated α-synuclein reveals that refolding of the unfolded PF is regulated by preNAC region.

Supplementary Figure 12. Effects of biochemical and thermal pre-treatments on the monomeric α-synuclein conformations.

Supplementary Figure 13. Illustrated diagram of the conformational structures and their transitions of monomeric α-synuclein.

References

#### Supplementary Methods

##### Protein expression and purification of α-synuclein-Ig27 fused proteins.

All the α-synuclein-Ig27 fused protein constructs were expressed in *E. Coli* DE3 bacteria with biotin protein ligase (BirA) and purified using the His-tag affinity column following the previous protocol[1]. Basically, the colonies were transfected with each of the corresponding plasmids and precultured in around 10 mL of LB medium containing 100 μg $mL^{-1}$ ampicillin at 37 °C overnight. The precultures were then inoculated into 1 L of ampicillin-containing LB medium and grown at 37 °C for around 4−6 hours until the optical density ($OD_{600}$) reached ~ 0.6. 50 µM biotin and 0.4 mM IPTG were then added into the cultures, and the resulting mixture was grown at 18 °C overnight. The biotin could be catalyzed by BirA ligase to conjugate to the AviTag peptide of the proteins in cell. Bacteria were harvested by centrifugation at 6000 g, and pellets were stored frozen at −80° C until further purification. In the purification steps, bacterial pellets were resuspended with lysis buffer (50 mM Tris, 300 mM NaCl, 1 mM PMSF, pH=7.4) and cells were mechanically lysed using French Press, followed by centrifugation at 40 000 g for 0.5 hours. The resultant supernatants were allowed to bind to $Co^{2+}$-NTA column (Thermo Scientific, DE, USA) at a cold room for 2 hours. After repeated washes (wash buffer: 50 mM Tris, 300 mM NaCl, 10 mM imidazole, pH=7.4), the proteins were eluted into elution buffer (50 mM Tris, 300 mM NaCl, 200 mM imidazole, pH=7.4). These proteins were further purified with gel filtration chromatography (Superdex 200, Äkta Pure system, GE Healthcare, MA, USA). The protein-containing fractions were verified using SDS–polyacrylamide gel electrophoresis, dialyzed into 1×PBS buffer, and frozen in aliquots with 15 % (v/v) glycerol by liquid nitrogen to be stored at −80 °C for use. Protein concentration was measured by spectrophotometry at 280 nm (NanoDrop 1000, Thermo Scientific, DE, USA).

**Protein expression and purification of α-synuclein monomer.** Full-length α-synuclein monomer (1−140) expression plasmid (pET21a-alpha-synuclein) was a gift from Michael J Fox Foundation MJFF (Addgene plasmid # 51486). The α-synuclein monomer was expressed and purified following the previous method[2,3]. Briefly, the bacterial induction started at an OD600 of ~ 0.6 with 0.4 mM IPTG for 4 hours at 37 °C. The harvested bacteria were resuspended into lysis buffer (50 mM Tris, 300 mM NaCl, 1 mM PMSF, 1mM EDTA, pH=7.4) and boiled in a water bath for 20 min at 100 °C. The solution was then centrifuged at 40 000 g for 30 minutes. The supernatant was dialyzed with 20 mM Tris (pH=7.0) overnight at 4 °C in SnakeSkin™ dialysis membrane (with a molecular weight cut off of 3 kDa, Thermo Fisher Scientific, USA). The α-synuclein monomer was then purified by ion exchange chromatography using Q-column (Hitrap Q FF column, GE Healthcare). The Q-column was firstly washed by Buffer B (1M NaCl 20 mM Tris, pH=7.0) and then equilibrated by Buffer A (20 mM Tris, pH=7.0). The α-synuclein solution was filtered through a 0.22 um filter, loaded onto the Q-column, washed by Buffer A buffer, and eluted against a linear gradient of ~ 10 column volumes of Buffer B using an ÄKTA Pure fast protein liquid chromatography (FPLC) system (GE Healthcare). The protein fractions were monitored on absorption at 280 nm and confirmed using SDS-PAGE. Fractions containing protein bands corresponding to the predicted monomer α-synuclein molecular weight (MW) of 14.4 kDa were further dialyzed overnight in 20 mM Tris pH 7.0 and concentrated with a 3 k Mw centrifugal concentrator to the desired concentration and stored at −80 °C.

**ThT fluorescence assay.** The purified α-synuclein-Ig27 fused protein or α-synuclein monomer (~100 μM) was diluted in 20 mM Tris buffer (pH=7.0) with 150 mM KCl, with or without seeds (~ 20 nM). The samples were mixed with 20 μM ThT and added to a 96-well plate. The plates were incubated at 37 °C with agitation (900 rpm) for 5 days. ThT signal was monitored using a microplate reader (Tecan) at an excitation wavelength of 440 nm and an emission wavelength of 490 nm. The resulting α-synuclein samples were characterized by Blue native PAGE (Novex) and AFM images.

**Blue native PAGE.** The native gel electrophoresis of α-synuclein samples is based on a Novex Bis-Tris Gel System using the XCell™ SureLock™ Mini-Cell (Life Technologies). In Blue Native PAGE, the Coomassie G-250 binds to proteins and confers a net negative charge while maintaining the proteins in their native state without any denaturation. The G-250 is present in the cathode buffer to provide a continuous flow of G-250 into the gel and is added to the α-synuclein samples containing non-ionic detergent before loading the samples onto the gel. The gel electrophoresis was running in an ice bath at 150 V for 2 hours. The gel was then washed and stained with Coomassie blue.

**AFM characterization.** AFM images of the α-synuclein samples were taken using a commercial AFM (Dimension FastScan, Icon Scanner, Bruker) in air under the tapping mode with a silicon cantilever (Fastscan A, Bruker). The samples were diluted and added to the mica substrate surfaces. After 10 min absorption, the mica substrate was washed by DI water and completely dried by blowing with nitrogen before imaging.

**Chamber and microbead preparation.** To measure the mechanical signature of the α-synuclein protein by MT setup, we prepared Spy-Catcher-functionalized membrane-well-based in-situ chambers and Neutravidin-coated magnetic beads for the formation of a single molecule tether through SpyTag−SpyCatcher attachment and biotin−Neutravidin cross-linking, respectively.

Superparamagnetic beads (Dynabeads™ M-270 Epoxy, 2.8 μm, Thermo Scientific) in dimethyl sulfoxide (DMSO) were diluted with 1 × PBS to a final concentration at around 1 mg $mL^{-1}$. The beads were pulled down by magnets and rinsed with 1 × PBS three times. 1 × PBS solution containing 50 μg $mL^{-1}$ (0.83 μM) neutravidin protein was added to the epoxy beads suspension. The mixture was kept away from light on a rotator and incubated at R.T. overnight, enabling the covalent bond formation of the reactive epoxy group to neutravidin protein. After 12 hours, the beads were washed repeatedly with 1 × PBS solution for ~ 5 times and blocked using BSA blocking buffer on a rotator for 12 hours. The neutravidin-coated magnetic beads were stored on a rotator at 4 °C for use.

Cover glasses (32 mm × 22 mm, 0.16 mm thick, Citoglass) were first immersed in detergent (50% Decon 90, Decon Laboratories Limited) overnight then sonicated for 30 minutes. After being thoroughly washed with deionized (DI) water, ethanol and acetone successively, these cover glasses were dried under steam of nitrogen and then treated with oxygen plasma (8% $O_2$, 50 mW, Tergeo plasma cleaner) for 5 minutes to expose surface hydroxyl groups. These glass substrates were immersed in an anhydrous methanol solution containing 1 % (v/v) APTES at room temperature (R.T.) for 1 hour. Then, they were washed sequentially using methanol and acetone, dried under a nitrogen flow, and incubated at 90 °C for 60 min. The fabrication of polymer microwells were prepared followed previous protocol[4]. Gently peel the polymer microwells prepared from slides and place on the middle of the bottom APTES-functionalized coverslip. Next, they were attached to a clean glass coverslip (18 mm × 18 mm, 0.10 mm thick, Cytoglass) using a SecureSeal adhesive sheet (0.12 mm thick, Sigma-Aldrich) to construct a in-situ laminar flow chamber. One entry of the channel was connected to a pump that can draw solution from the channel to a syringe, while the other entry was connected with a tube for flowing in new solution. This chamber was then immersed with Milli-Q water and buffer-exchanged to 1% (v/v) glutaraldehyde solution. After the two-hour glutaraldehyde treatment, the chamber was gently washed with water followed by 1 × PBS solution. About 100 μL of 1 × PBS solution containing polystyrene beads (1:100 dilution, 2.68 μm amino polystyrene beads, Spherotech) has flowed into the chamber. The polystyrene beads were used as references adhered on the surface to eliminate spatial drift during experiments. After incubating for 1 hour, 1 × PBS solution with 25 μg $mL^{-1}$ (2 μM) spy-catcher protein has seeped into the chamber. The chamber was kept wet and allowed to incubate at room temperature (R.T.) for 4 hours, allowing for the conjugation of spy-catcher protein to the cover glass. Unconjugated proteins were then washed away with 1 × PBS a few times. Finally, the chamber was treated with blocking buffer (1 × PBS, 3% BSA, 0.02% $NaN_3$, pH=7.4) at R.T. overnight to prevent non-specific binding in the following experiments. The chambers were kept wet and stored at 4 °C until use.

**Sample preparation.** For the wild-type α-synuclein and its truncated variants, the proteins were taken from −80 °C stocks and diluted to approximately 1 nM using the standard working buffer (20 mM HEPES, 50 mM KCl, 10 mM $MgCl_2$, 1% (m/m) BSA, 1 mM DTT, pH = 7.4) for sample preparation. The spy-catcher-functionalized chamber was first washed for a few times with the standard buffer, then the diluted proteins containing spy-tag were introduced into the chamber and incubated for 30 minutes, allowing the surface anchorage to occur between the recombinant protein and the chamber surface via spy-tag−spy-catcher interaction. The chamber was washed a few times to remove the unbounded proteins. Next, an adequate amount of Neutravidin-coated superparamagnetic beads was added into the chamber and allowed to incubate for 15 minutes. Finally, the chamber was sealed completely using mineral oil to mitigate the effects of solution evaporation during the long-time experiment. The chamber was then transferred and mounted onto the magnetic-tweezer setup to proceed with the single-molecule experiment.

**Magnetic tweezer setup and single-molecule force scan.** The homemade magnetic tweezer was built on an inverted microscope (IX71, Olympus). 100 × oil-immersion objective (UPlanFLN 100X, Olympus) was used to monitor images of the protein-tethered bead. An image library at a series of different focal planes was built by adjusting the focal plane of the objective with a piezo objective actuator (E-753 Digital Piezo Controller, PI). Two permanent magnetic rods were placed vertically above the chamber, whose movement was controlled by a motorized stage (VT-40, Micos) to change the magnetic force applied to the sample. By adopting a high FOV camera (Basler acA1920-155 μm), multiple magnetic beads could be tracked and monitored in real time. During the measurements, the height changes of the magnetic beads compared to the fixed reference beads, which corresponded to the extension of the protein tether, were recorded. The piezo objective actuator was used to correct the drift of the focal plane in real-time by monitoring the drift of reference bead, which provided a long-term ultra stability of the magnetic tweezer setup.

In a typical single-molecule force scan experiment, the magnet was moved from approximately 1.5 pN to around 100 pN at a constant loading rate in the range of 0.5−5.0 pN $s^{-1}$. Subsequently, the force applied to the tethered bead was decreased to around 1.5 pN at a constant loading rate in the range of -0.2 − -2.0 pN $s^{-1}$. After recovery at 1.5 pN for a time duration of 30 s, the tethered single molecule was scanned for the next round by repeating the force increasing and decreasing cycles. The bead height-time traces of the tethers were recorded in real-time. Collectively, by using this magnetic tweezer setup, we were able to record force scan data over a period from hours to days for multiple tethers.

**Pre-treatment experiments**

In the pre-treatment experiments, α-synuclein proteins were subjected to direct treatment with different agents: (1) 2% SDS in 1 × PBS (pH=7.4) buffer for 1 hour; (2) denaturant including 8 M urea and 6 M GuHCl in 1 × PBS (pH=7.4) for 1 hour. In the recovering experiment, α-synuclein proteins were first treated with 8 M urea and 6 M GuHCl in 1 × PBS (pH=7.4) buffer for 1 hour. The completely denatured proteins were then buffer-exchanged using an ultra-spin column to remove the denaturant and incubated: (1) at 23 ºC or 37 ºC for 5 days, with aliquoted fractions taken at 6 h, 12 h, 24 h, 48 h, 72 h, and 120 h for single-molecule assays; (2) with 1 µM α-synuclein aggregates (seed) at 23 ºC for 24 h. In the pre-treatment experiment using small-molecule drugs, α-synuclein proteins were directly treated with 100 µM of each of the following compounds: EGCG, Alne 138b, fasudil, curcumin, bailcalein, and dopamine in 20 mM HEPES, 50 mM KCl, 10 mM $MgCl_2$, pH=7.4 for 24 hours at 23 ºC. After these biochemical or drug perturbations, α-synuclein proteins were diluted to approximately 1 nM in the standard working buffer for single-molecule experiments.

**Force calibration**

Force calibration was conducted for each tethered bead measured in the experiments, which had ~10 % relative error[5,6]. The force applied to a bead is a function of the distance ($d$) between permanent magnets and the paramagnetic beads, which was calibrated by 16 µm λ-DNA as described in our previous publication[5]. For the magnets pair we used in the manuscript, the force equation is given by[5]:

$$f = C * \left(exp\left(-\frac{d}{0.36}\right) + 0.48\, exp\left(-\frac{d}{1.12}\right)\right) \qquad \text{Eq. 1}$$

where $d$ is in the unit of millimeter, and $C$ value is a constant which differs for each paramagnetic bead. For the M270 beads, the $C$ value is in the range of 180−210 pN.

For each individual tethered bead measured in experiments, the force was calibrated by the bead fluctuation at forces < 10 pN by[5,6]:

$$f = \frac{k_B T(R+z)}{{\delta_y}^2} \qquad \text{Eq. 2}$$

where $k_B T$ is the Boltzmann constant times temperature, $R$ is the radius of the paramagnetic bead (1.4 µm for typical M270 Dynabead), $z$ is the extension of tether, and ${\delta_y}^2$ is the variance of transverse fluctuation of the magnetic bead perpendicular to the magnetization direction. The calibrated forces were extrapolated to higher forces through Eq. 1. Due to the uncertainty of bead size and the deviation of the tether point from the bottom pole of the bead, the calibrated forces have about 10% relative error.

**Protein construction and sequence**

The full amino acid sequences of the protein construction used in this work are listed below. All sequences contain a 6×HIS (HHHHHH) tag for purification, an avi-tag (GLNDIFEAQKIEWHE) for biotinylation, and a spy-tag (AHIVMVDAYKPTK) for surface covalent functionalization.

**α-synuclein-WT-I27:** avi-2I27-α-synuclein-WT(1-140)-2I27-spy

MHHHHHHGKPIPNPLLGLDSTENLYFQGIDPFTGLNDIFEAQKIEWHEGGGSGLI
EVEKPLYGVEVFVGETAHFEIELSEPDVHGQWKLKGQPLAASPDAEIIEDGKKHILIL
HNAQLGMTGEVSFQAANTKSAANLKVKELGGGSGLIEVEKPLYGVEVFVGETAHFE
IELSEPDVHGQWKLKGQPLAASPDAEIIEDGKKHILILHNAQLGMTGEVSFQAANTKS
AANLKVKELGGGSGKLGGGSGMDVFMKGLSKAKEGVVAAAEKTKQGVAEAAGKT

KEGVLYVGSKTKEGVVHGVATVAEKTKEQVTNVGGAVVTGVTAVAQKTVEGAGSI
AAATGFVKKDQLGKNEEGAPQEGILEDMPVDPDNEAYEMPSEEGYQDYEPEAGGG
SGLEGGGSGLIEVEKPLYGVEVFVGETAHFEIELSEPDVHGQWKLKGQPLAASPDAEI
IEDGKKHILILHNAQLGMTGEVSFQAANTKSAANLKVKELGGGSGLIEVEKPLYGVE
VFVGETAHFEIELSEPDVHGQWKLKGQPLAASPDAEIIEDGKKHILILHNAQLGMTGE
VSFQAANTKSAANLKVKELGGGSGAHIVMVDAYKPTK

**α-synuclein-NAC-I27:** avi-2I27-α-synuclein-NAC(61-100)-2I27-spy

MHHHHHHGKPIPNPLLGLDSTENLYFQGIDPFTGLNDIFEAQKIEWHEGGGSGLI
EVEKPLYGVEVFVGETAHFEIELSEPDVHGQWKLKGQPLAASPDAEIIEDGKKHILIL
HNAQLGMTGEVSFQAANTKSAANLKVKELGGGSGLIEVEKPLYGVEVFVGETAHFE
IELSEPDVHGQWKLKGQPLAASPDAEIIEDGKKHILILHNAQLGMTGEVSFQAANTKS
AANLKVKELGGGSGKLGGGSGEQVTNVGGAVVTGVTAVAQKTVEGAGSIAAATGF
VKKDQLGGGSGLEGGGSGLIEVEKPLYGVEVFVGETAHFEIELSEPDVHGQWKLKG
QPLAASPDAEIIEDGKKHILILHNAQLGMTGEVSFQAANTKSAANLKVKELGGGSGLI
EVEKPLYGVEVFVGETAHFEIELSEPDVHGQWKLKGQPLAASPDAEIIEDGKKHILIL
HNAQLGMTGEVSFQAANTKSAANLKVKELGGGSGAHIVMVDAYKPTK

**α-synuclein-NAC+preNAC-I27:** avi-2I27-α-synuclein-NAC+preNAC(36-100)-2I27-spy

MHHHHHHGKPIPNPLLGLDSTENLYFQGIDPFTGLNDIFEAQKIEWHEGGGSGLI
EVEKPLYGVEVFVGETAHFEIELSEPDVHGQWKLKGQPLAASPDAEIIEDGKKHILIL
HNAQLGMTGEVSFQAANTKSAANLKVKELGGGSGLIEVEKPLYGVEVFVGETAHFE
IELSEPDVHGQWKLKGQPLAASPDAEIIEDGKKHILILHNAQLGMTGEVSFQAANTKS
AANLKVKELGGGSGKLGGGSGGVLYVGSKTKEGVVHGVATVAEKTKEQVTNVGG
AVVTGVTAVAQKTVEGAGSIAAATGFVKKDQLGGGSGLEGGGSGLIEVEKPLYGVE
VFVGETAHFEIELSEPDVHGQWKLKGQPLAASPDAEIIEDGKKHILILHNAQLGMTGE
VSFQAANTKSAANLKVKELGGGSGLIEVEKPLYGVEVFVGETAHFEIELSEPDVHGQ
WKLKGQPLAASPDAEIIEDGKKHILILHNAQLGMTGEVSFQAANTKSAANLKVKELG
GGSGAHIVMVDAYKPTK

**α-synuclein-ΔC-I27:** avi-2I27-α-synuclein-ΔC(1-100)-2I27-spy

MHHHHHHGKPIPNPLLGLDSTENLYFQGIDPFTGLNDIFEAQKIEWHEGGGSGLI
EVEKPLYGVEVFVGETAHFEIELSEPDVHGQWKLKGQPLAASPDAEIIEDGKKHILIL
HNAQLGMTGEVSFQAANTKSAANLKVKELGGGSGLIEVEKPLYGVEVFVGETAHFE
IELSEPDVHGQWKLKGQPLAASPDAEIIEDGKKHILILHNAQLGMTGEVSFQAANTKS
AANLKVKELGGGSGKLGGGSGMDVFMKGLSKAKEGVVAAAEKTKQGVAEAAGKT
KEGVLYVGSKTKEGVVHGVATVAEKTKEQVTNVGGAVVTGVTAVAQKTVEGAGSI
AAATGFVKKDQLGGGSGLEGGGSGLIEVEKPLYGVEVFVGETAHFEIELSEPDVHGQ
WKLKGQPLAASPDAEIIEDGKKHILILHNAQLGMTGEVSFQAANTKSAANLKVKELG
GGSGLIEVEKPLYGVEVFVGETAHFEIELSEPDVHGQWKLKGQPLAASPDAEIIEDGK
KHILILHNAQLGMTGEVSFQAANTKSAANLKVKELGGGSGAHIVMVDAYKPTK

**α-synuclein-ΔN1-I27:** avi-2I27-α-synuclein-ΔN1(11-140)-2I27-spy

MHHHHHHGKPIPNPLLGLDSTENLYFQGIDPFTGLNDIFEAQKIEWHEGGGSGLI
EVEKPLYGVEVFVGETAHFEIELSEPDVHGQWKLKGQPLAASPDAEIIEDGKKHILIL
HNAQLGMTGEVSFQAANTKSAANLKVKELGGGSGLIEVEKPLYGVEVFVGETAHFE

IELSEPDVHGQWKLKGQPLAASPDAEIIEDGKKHILILHNAQLGMTGEVSFQAANTKS AANLKVKELGGGSGKLGGGSGAKEGVVAAAEKTKQGVAEAAGKTKEGVLYVGSK TKEGVVHGVATVAEKTKEQVTNVGGAVVTGVTAVAQKTVEGAGSIAAATGFVKK DQLGKNEEGAPQEGILEDMPVDPDNEAYEMPSEEGYQDYEPEAGGGSGLEGGGSGL IEVEKPLYGVEVFVGETAHFEIELSEPDVHGQWKLKGQPLAASPDAEIIEDGKKHILIL HNAQLGMTGEVSFQAANTKSAANLKVKELGGGSGLIEVEKPLYGVEVFVGETAHFE IELSEPDVHGQWKLKGQPLAASPDAEIIEDGKKHILILHNAQLGMTGEVSFQAANTKS AANLKVKELGGGSGAHIVMVDAYKPTK

**α-synuclein-ΔN2-I27:** avi-2I27-α-synuclein-ΔN2(36-140)-2I27-spy

MHHHHHHGKPIPNPLLGLDSTENLYFQGIDPFTGLNDIFEAQKIEWHEGGGSGLI EVEKPLYGVEVFVGETAHFEIELSEPDVHGQWKLKGQPLAASPDAEIIEDGKKHILIL HNAQLGMTGEVSFQAANTKSAANLKVKELGGGSGLIEVEKPLYGVEVFVGETAHFE IELSEPDVHGQWKLKGQPLAASPDAEIIEDGKKHILILHNAQLGMTGEVSFQAANTKS AANLKVKELGGGSGKLGGGSGGVLYVGSKTKEGVVHGVATVAEKTKEQVTNVGG AVVTGVTAVAQKTVEGAGSIAAATGFVKKDQLGKNEEGAPQEGILEDMPVDPDNE AYEMPSEEGYQDYEPEAGGGSGLEGGGSGLIEVEKPLYGVEVFVGETAHFEIELSEP DVHGQWKLKGQPLAASPDAEIIEDGKKHILILHNAQLGMTGEVSFQAANTKSAANL KVKELGGGSGLIEVEKPLYGVEVFVGETAHFEIELSEPDVHGQWKLKGQPLAASPDA EIIEDGKKHILILHNAQLGMTGEVSFQAANTKSAANLKVKELGGGSGAHIVMVDAY KPTK

**α-synuclein-ΔN3-I27:** avi-2I27-α-synuclein-ΔN3(61-140)-2I27-spy

MHHHHHHGKPIPNPLLGLDSTENLYFQGIDPFTGLNDIFEAQKIEWHEGGGSGLI EVEKPLYGVEVFVGETAHFEIELSEPDVHGQWKLKGQPLAASPDAEIIEDGKKHILIL HNAQLGMTGEVSFQAANTKSAANLKVKELGGGSGLIEVEKPLYGVEVFVGETAHFE IELSEPDVHGQWKLKGQPLAASPDAEIIEDGKKHILILHNAQLGMTGEVSFQAANTKS AANLKVKELGGGSGKLGGGSGEQVTNVGGAVVTGVTAVAQKTVEGAGSIAAATGF VKKDQLGKNEEGAPQEGILEDMPVDPDNEAYEMPSEEGYQDYEPEAGGGSGLEGG GSGLIEVEKPLYGVEVFVGETAHFEIELSEPDVHGQWKLKGQPLAASPDAEIIEDGKK HILILHNAQLGMTGEVSFQAANTKSAANLKVKELGGGSGLIEVEKPLYGVEVFVGET AHFEIELSEPDVHGQWKLKGQPLAASPDAEIIEDGKKHILILHNAQLGMTGEVSFQAA NTKSAANLKVKELGGGSGAHIVMVDAYKPTK

**α-synuclein-ΔNAC-I27:** avi-2I27-α-synuclein-ΔNAC(Δ61-100)-2I27-spy

MHHHHHHGKPIPNPLLGLDSTENLYFQGIDPFTGLNDIFEAQKIEWHEGGGSGLI EVEKPLYGVEVFVGETAHFEIELSEPDVHGQWKLKGQPLAASPDAEIIEDGKKHILIL HNAQLGMTGEVSFQAANTKSAANLKVKELGGGSGLIEVEKPLYGVEVFVGETAHFE IELSEPDVHGQWKLKGQPLAASPDAEIIEDGKKHILILHNAQLGMTGEVSFQAANTKS AANLKVKELGGGSGKLGGGSGMDVFMKGLSKAKEGVVAAAEKTKQGVAEAAGKT KEGVLYVGSKTKEGVVHGVATVAEKTKGKNEEGAPQEGILEDMPVDPDNEAYEMP SEEGYQDYEPEAGGGSGLEGGGSGLIEVEKPLYGVEVFVGETAHFEIELSEPDVHGQ WKLKGQPLAASPDAEIIEDGKKHILILHNAQLGMTGEVSFQAANTKSAANLKVKELG GGSGLIEVEKPLYGVEVFVGETAHFEIELSEPDVHGQWKLKGQPLAASPDAEIIEDGK KHILILHNAQLGMTGEVSFQAANTKSAANLKVKELGGGSGAHIVMVDAYKPTK

**α-synuclein-ΔpreNAC-I27:** avi-2I27-α-synuclein-ΔpreNAC(Δ36-60)-2I27-spy

MHHHHHHGKPIPNPLLGLDSTENLYFQGIDPFTGLNDIFEAQKIEWHEGGGSGLI
EVEKPLYGVEVFVGETAHFEIELSEPDVHGQWKLKGQPLAASPDAEIIEDGKKHILIL
HNAQLGMTGEVSFQAANTKSAANLKVKELGGGSGLIEVEKPLYGVEVFVGETAHFE
IELSEPDVHGQWKLKGQPLAASPDAEIIEDGKKHILILHNAQLGMTGEVSFQAANTKS
AANLKVKELGGGSGKLGGGSGMDVFMKGLSKAKEGVVAAAEKTKQGVAEAAGKT
KEEQVTNVGGAVVTGVTAVAQKTVEGAGSIAAATGFVKKDQLGKNEEGAPQEGILE
DMPVDPDNEAYEMPSEEGYQDYEPEAGGGSGLEGGGSGLIEVEKPLYGVEVFVGET
AHFEIELSEPDVHGQWKLKGQPLAASPDAEIIEDGKKHILILHNAQLGMTGEVSFQAA
NTKSAANLKVKELGGGSGLIEVEKPLYGVEVFVGETAHFEIELSEPDVHGQWKLKG
QPLAASPDAEIIEDGKKHILILHNAQLGMTGEVSFQAANTKSAANLKVKELGGGSGA
HIVMVDAYKPTK

**α-synuclein-monomer:** α-synuclein-WT(1-140) (PET21a)

MDVFMKGLSKAKEGVVAAAEKTKQGVAEAAGKTKEGVLYVGSKTKEGVVHG
VATVAEKTKEQVTNVGGAVVTGVTAVAQKTVEGAGSIAAATGFVKKDQLGKNEEG
APQEGILEDMPVDPDNEAYEMPSEEGYQDYEPEA

**α-synuclein-WT:** avi-α-synuclein-WT(1-140)-spy (PET21a)

MGLNDIFEAQKIEWHEGGGSGKLGGGSGMDVFMKGLSKAKEGVVAAAEKTK
QGVAEAAGKTKEGVLYVGSKTKEGVVHGVATVAEKTKEQVTNVGGAVVTGVTAV
AQKTVEGAGSIAAATGFVKKDQLGKNEEGAPQEGILEDMPVDPDNEAYEMPSEEGY
QDYEPEAGGGSGLEGGGSGAHIVMVDAYKPTK

### Supplementary Notes

#### 1. The role of I27 domains in the recombinant protein

Fusing well-characterized protein domains into the recombinant protein construction of the single-molecule study is helpful to identify the unknown mechanical responses of the protein to be studied, which is a widely used strategy in single-molecule studies[7,8].

All experiments using the I27-fused-α-synuclein in the manuscript were obtained from the single-molecule tethers with the highly characteristic unfolding signals of the four I27 domains[9]. To confirm that the observed mechanical signatures are not affected by the I27 domains, we also performed control experiments using the α-synuclein construct without the I27 domain. For a such construct, the Avi-α-synuclein-Spytag protein was tethered between a Neutravidin-coated superparamagnetic bead and a SpyCatcher-coated coverslip glass surface via a 576 bp dsDNA spacer as illustrated in Supplementary Fig. 5A. Single, monomeric tether is ensured based on the unique overstretching transition of dsDNA that occurs slightly above 65 pN forces[10]. The characteristic unfolding and refolding signals of PF α-synuclein were still observed in such construct without the I27 domains (Supplementary Fig. 5B−C). These results confirmed that the observed PF intermediate conformation was from the tethered monomeric α-synuclein.

We further studied the aggregation behavior of the recombinant protein constructs in bulk testing. Unlike the α-synuclein monomer or I27-truncated α-synuclein, we found that the recombinant α-synuclein protein fused with four I27 domains did not form higher-order aggregates or oligomers (Supplementary Fig. 1). Similar results were previously reported using MBP domains fused α-synuclein[11], where the fused protein domains inhibit the aggregation of α-synuclein. In another work using I27-fused α-synuclein, they show that the α-synuclein did undergo an α-to-β structural transition during the incubation using CD spectra[12], which is related to the aggregation precursor of synucleinopathy. Together, these results indicate that the four I27 tandem repeats along the N- and C- terminal of α-synuclein did not bias the monomeric behavior of the α-synuclein, but worked as a protector avoiding any aggregation or oligomerization of the recombinant protein because of the steric effect. In this way, the recombinant α-synuclein protein then served as a model system to study the monomeric behavior of α-synuclein.

#### 2. The observed smooth force-bead height curves

There are two potential explanations for the observed smooth force-bead height curves: either they indicate a completely random, unstructured coiled conformation of α-synuclein, or they suggest a conformation that might include certain ultra-stable compact structures that did not unfold during the force-loading experiments. The latter possibility is highly improbable, given the absence of a stepwise increase in bead height throughout 50 hours of repeated force scanning experiments or at our highest constant forces attainable (115 ± 11.5 pN) for over 10 hours. Moreover, the stepwise increase in bead height was also absent following the in-situ introduction of a detergent (GuHCl/urea) that expected to completely denature a protein's secondary structure (Supplementary Fig. 3).

#### 3. Force jump experiments of the PF α-synuclein

Apart from the force scanning experiment with constant force increasing or force decreasing loading rates, we also performed force jump experiments to measure the unfolding/refolding lifetimes at constant forces of PF α-synuclein. The representative bead height−time traces containing unfolding and refolding events of PF α-synuclein in force-jumping cycles are shown in Supplementary Fig. 6A. The applied external force is jumping from a resting force of 1.5 ± 0.15 pN to an unfolding force of 56.5 ± 5.7 pN, which is to allow the unfolding of the PF intermediate, and then jumping to a refolding force of 10.6 ± 1.1 pN, which is to allow the refolding of the PF intermediate. By repeating such force-jumping operations on multiple molecules for multiple cycles, the force-dependent unfolding and refolding lifetime of the PF intermediate at each force can be obtained. The corresponding unfolding (at 56.5 ± 5.7 pN) and refolding lifetime histograms (10.6 ± 1.1 pN) are shown in Supplementary Fig. 6B, which are well fitted by a single exponential decay to obtain the average lifetime $\tau_f$ at a giving mechanical force. The average force-dependent transition rates of unfolding and refolding of PF α-synuclein are shown in Supplementary Fig. 6C.

#### 4. The minor conformational species

We have observed a minor fraction (~5%) of the α-synuclein species with distinct mechanical signatures compared to the PF. Supplementary Fig. 7A shows a representative force-height trace of a typical minor conformational species in the force scan, which is characterized by a two-stepwise bead height increases at unfolding forces at ~8 pN and ~15 pN, respectively (Supplementary Fig. 7B). The two unfolding step sizes are in the ranges of 12−15 nm and 15−20 nm, respectively (Supplementary Fig. 7B), suggesting the existence of two partially folded intermediate structures. For the two dynamic intermediates, rapid (seconds time scale) reversible transitions between the folded and unfolded states were observed at a constant force at around 6.2 pN (Supplementary Fig. 7C).

#### 5. The pre-treatment experiments

In order to ensure identical initial conditions for the investigations, the monomeric α-synuclein proteins were fully unfolded using 6 molars (M) guanidine hydrochloride (GuHCl) and 8 M urea, and subsequently recovered under different biochemical conditions following the removal of the denaturants, which ensure identical initial conditions for all the subsequent recovery experiments. After pre-treatment, the fraction of conformational species of the monomeric α-synuclein was investigated using the single-molecule loading rate scanning experiments (Supplementary Fig. 12a).

The conformational fractions of α-synuclein resulting from the applied perturbations in pre-treatment experiments are presented in Supplementary Fig. 12b and compared to the untreated control. Treatment with 6 M GuHCl and 8 M urea resulted in a dramatically decreased PF fraction to a negligible level, leaving the disordered conformation as the predominant species. This finding is consistent with the ability of the selected denaturants (GuHCl and urea) to fully convert monomeric α-synuclein into an unstructured conformation, regardless of its initial state following protein expression and purification. In contrast, sodium dodecyl sulphate (SDS), a denaturant previously shown to induce α-synuclein fibrillation[13], did not reduce the occurrence of PF (Supplementary Fig. 12b).

We then investigated the impact of an aggregation-promoting prion-like seed, comprising small fibrils of α-synuclein. Following removal of the denaturants and a 24-hour incubation with seed, the fraction of the PF species escalated to approximately 50 % in the presence of 1 μM seed (equivalent to the molar concentration of monomer) at 23 °C. In stark contrast, in the absence of the seed, incubation over the same duration resulted in a considerably lower level of PF species (~ 10%) at the same temperature. This result suggests that the seed, recognized for its ability to trigger α-synuclein fibril formation[14,15], also promotes the generation of the PF intermediate.

We next investigated the fractions of different α-synuclein conformations after incubation at two temperatures, 23 °C or 37 °C, over various time durations up to 120 hours. As shown in Supplementary Fig. 12c, at both temperatures, the PF fraction increased with prolonged incubation time, with a faster increase observed at 37 °C compared to 23 °C. Specifically, after 24 hours incubation, approximately 40% of α-synuclein exists in the PF fraction at 37 °C, whereas only around 10% of the PF fraction was observed at 23 °C (Supplementary Fig. 12b−c). These results suggest that a higher temperature facilitates the formation of the PF species in monomeric α-synuclein. Interestingly, higher temperatures have also been shown to increase the rate of higher-order aggregation of α-synuclein[16,17], making this finding particularly noteworthy.

#### Supplementary Figures

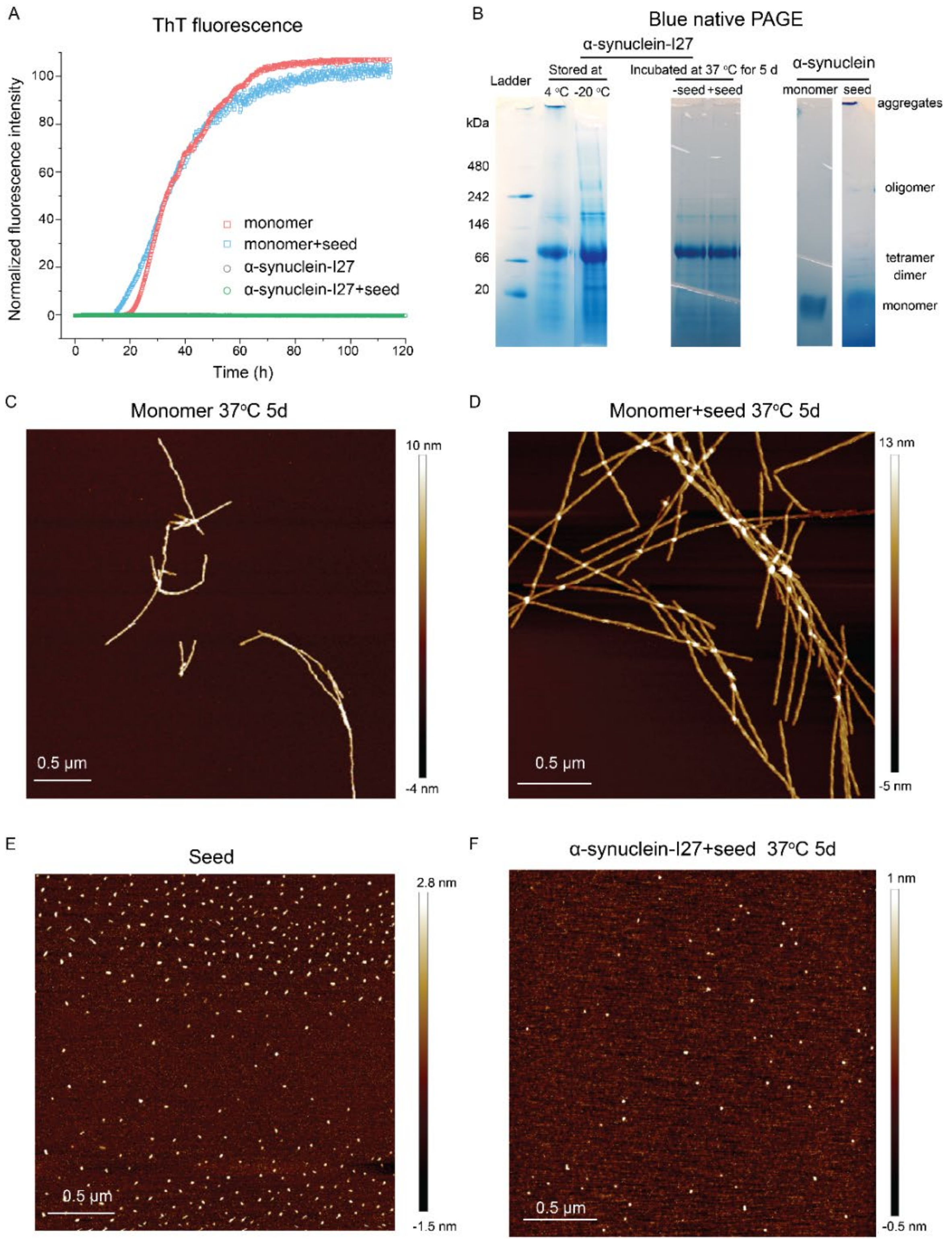


**Supplementary Figure 1. Characterizations of ThT fluorescence, AFM image, and native gel of the α-synuclein-I27 and α-synuclein monomer samples.** (A) ThT fluorescence spectra reveal the aggregation kinetics of 100 μM α-synuclein-I27 and α-synuclein monomer. The proteins were incubated with/without 20 nM seeds (small α-synuclein aggregates) in 20 mM Tris, 150 mM KCl, pH=7.0 at 37 °C with agitation at 600 rpm and measured at least in triplicate. The ThT fluorescence signals were converted to normalized intensities. (B) Blue native PAGE of the α-synuclein samples (from left to right): ladders, fresh wild-type α-synuclein-I27 proteins stored at 4 ºC or −20 ºC, wild-type α-synuclein-I27 proteins incubated at 37 ºC for 5 days with/without 20 nM seeds, α-synuclein monomer, α-synuclein seeds. (C-F) AFM characterization of the α-synuclein samples: (C) α-synuclein monomer incubated at 37 ºC for 5 days, (D) α-synuclein monomer incubated with 20 nM seeds at 37 ºC for 5 days, (E) α-synuclein seeds (ultrasonicated α-synuclein fibrils), (F) α-synuclein-I27 proteins with 20 nM seeds incubated at 37 ºC for 5 days.

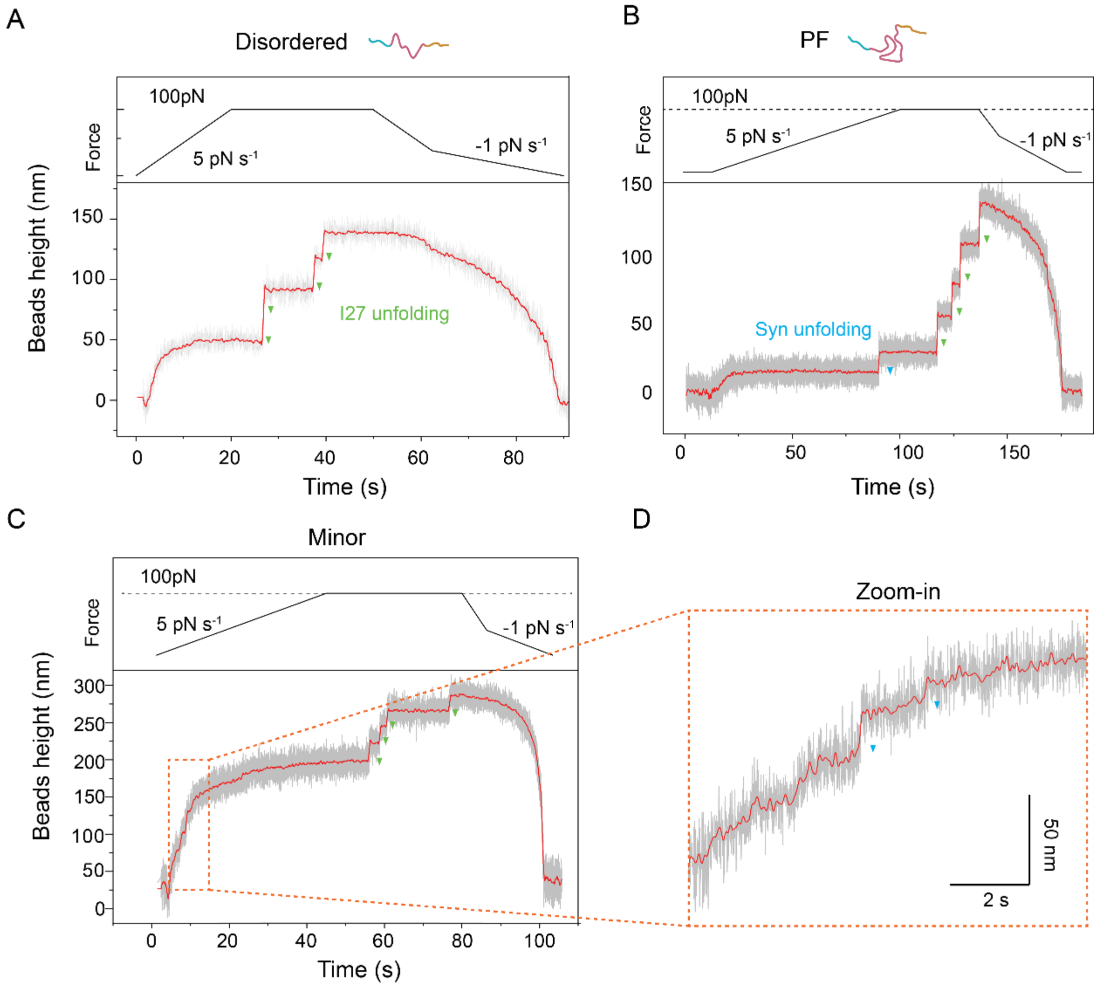


**Supplementary Figure 2. The mechanical responses of wild-type monomeric α-synuclein-I27-fused protein.** (A) Representative force–time trace of disordered α-synuclein-I27 protein, where only the characteristic unfolding of the four I27 domains is observed. (B) Representative force–time trace of α-synuclein-I27 protein with the mechanical stable partially folded (PF) intermediate. (C) Representative force–time trace of α-synuclein-I27 protein in the minor conformation. (D) Enlarged force-time trace from (C) reveals two-stepwise jumps during the force increasing scan. The forces are increased from ~1.5 pN to ~ 100 pN at a constant loading rate of 5 pN $s^{-1}$and subsequently held at 100 pN until the four I27 unfolding. The forces were then rapidly decreased to ~ 10 pN at a constant loading rate of −5 pN $s^{-1}$ and subsequently decreased to ~ 1.5 pN at a slower force loading rate of −1 pN $s^{-1}$ to allow the proper refolding of the proteins. The raw data and 200-FFT smoothed data are shown in grey and red, respectively. Green triangles represent the unfolding steps of I27, while blue triangles represent the unfolding signal of α-synuclein.

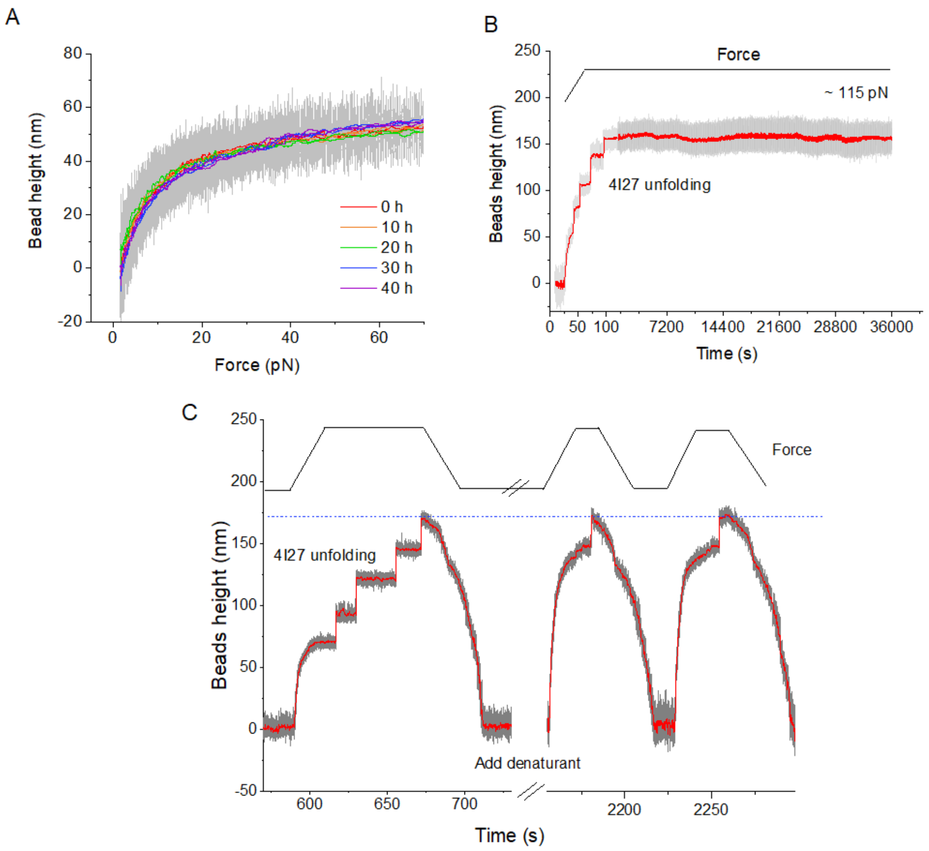


**Supplementary Figure 3. The mechanical responses of disordered α-synuclein.** (A) Force-bead height curves recorded at 5 pN $s^{-1}$ of a disordered α-synuclein over 50 hours of repeating force scan experiments, where no I27 unfolding occurs within this force range at the loading rate[9]. (B) Bead height−time trace of a disordered α-synuclein at a high constant force (~115 pN) for more than 10 hours. Only unfolding of the four I27 domains was observed. (C) Bead height−time trace of a disordered α-synuclein under in-situ treatment of denaturant (6 M GuHCl and 8 M urea solution) to disrupt the secondary structure of the proteins. The representative data show three of four I27 domains were unfolded by the treatment over the time scale. Similar results were observed at three different tethered proteins.

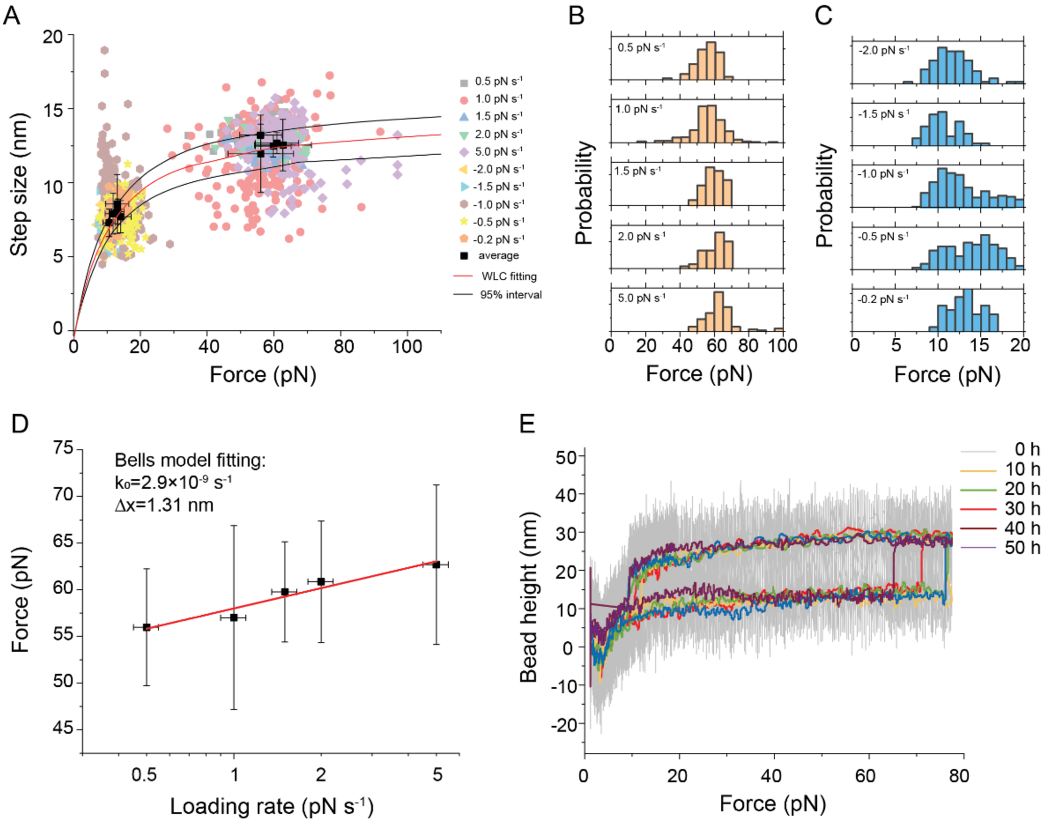


**Supplementary Figure 4. The mechanical responses of PF α-synuclein under loading rate force scan.** (A) Step sizes of the unfolding and refolding of the partially folded intermediate in PF α-synuclein observed in the force scan at different force-increasing or decreasing loading rates, which were collected from more than 30 different tethered PF α-synuclein. Raw data are presented as transparent colored scatters. The numbers of observed unfolding events are: 84, 225, 68, 47, and 124 for loading rate of 0.5 pN $s^{-1}$, 1.0 pN $s^{-1}$, 1.5 pN $s^{-1}$, 2.0 pN $s^{-1}$, and 5.0 pN $s^{-1}$, respectively. The numbers of observed refolding events are: 105, 59, 268, 126, and 44 for loading rate of -2.0 pN $s^{-1}$, -1.5 pN $s^{-1}$, -1.0 pN $s^{-1}$, -0.5 pN $s^{-1}$, and -0.2 pN $s^{-1}$, respectively. The black solid squares denote the average step size, average unfolding force, and average refolding force at each loading rate. Error bar indicates S.D.. A Worm-like chain model was used to fit the data (red line), and the optimal fitting parameters were obtained as A= 0.50 ± 0.05 nm, Lc= 15.50 ± 1.50 nm. Considering an average contour length of 0.38 nm[18] per amino acid, the number of amino acids involved in the partially folded structure is about 41 (a.a.). (B) The unfolding force distribution of the PF α-synuclein at different force-increasing loading rates. (C) The refolding force distribution of the at different force-decreasing loading rates. (D) Dynamic force spectra show the average unfolding forces of the PF α-synuclein at different loading rates. Red solid line is the best fitting to the force-loading rate using the Bell-Evans model ( $\boldsymbol{F(r)} = \frac{\boldsymbol{k_B T}}{\boldsymbol{\Delta x}} \mathbf{exp}\,(\frac{\boldsymbol{r\Delta x}}{\boldsymbol{k_0 k_B T}})$ )[19,20], where the dissociate rate constant $\boldsymbol{k_0} = (\mathbf{2.9} \pm \mathbf{7.69}) \times \mathbf{10^{-9}}$ $s^{-1}$ and transition distance $\boldsymbol{\Delta x} = \mathbf{1.30} \pm \mathbf{0.19}$ nm. ± indicates S.E.. Error bars indicate the mean ± S.D.. (E) Force-extension curves of a PF α-synuclein over 50 hours of repeating force scanning experiments.

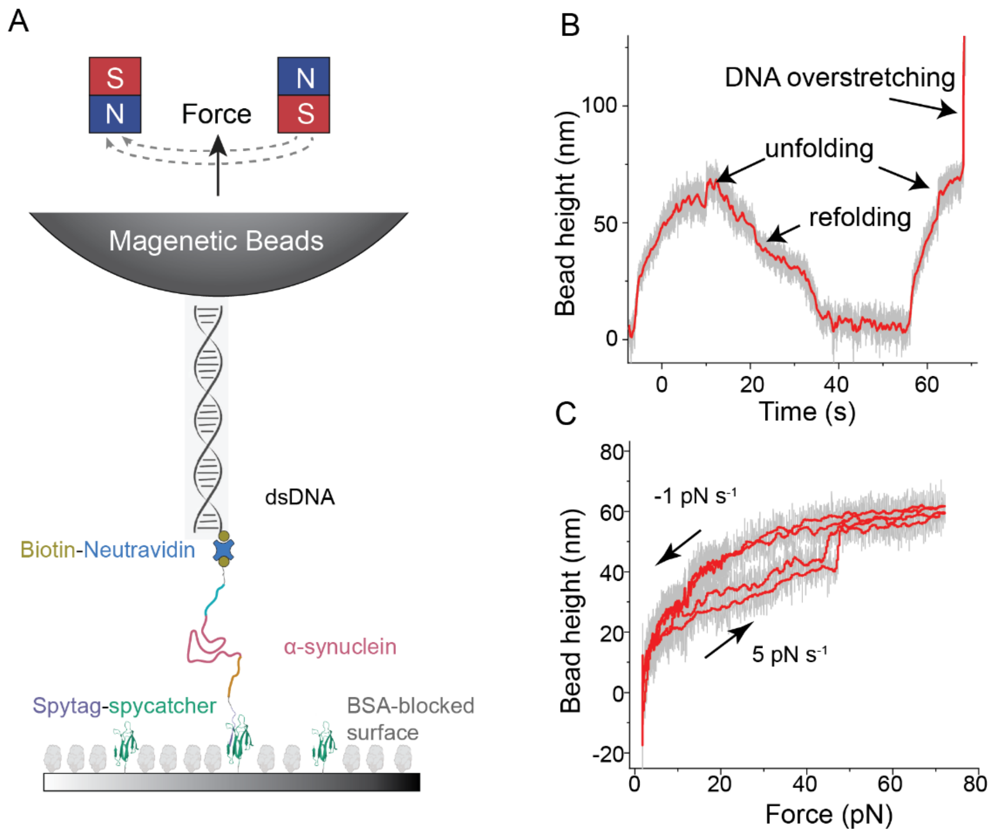


**Supplementary Figure 5. Control experiments using the $\alpha$-synuclein protein without the I27 domains.** (A) The schematic diagram of the single-molecule magnetic-tweezer experiment using the Avi-α-synuclein-Spy protein construct with a double-strand DNA (dsDNA) handle spacer. The recombinant monomeric $\alpha$-synuclein was tethered to the superparamagnetic beads (Dynabeads M270) via a 576bp-dsDNA handle. (B) Typical bead height time traces and (C) force-extension curves of a PF monomeric $\alpha$-synuclein in force loading rate scan, where the stretching force is increased at a constant loading rate of 5 pN $s^{-1}$ and subsequently decreased to around 1.5 pN at a constant loading rate of -1 pN $s^{-1}$. The overstretching extension of dsDNA at ~65 pN could be used as a fingerprint. Raw data and smoothed curves are indicated in grey and red, respectively. We note that due to the overstretching extension of dsDNA at ~65 pN, the bead height-force traces had a cut-off at around 60 pN.

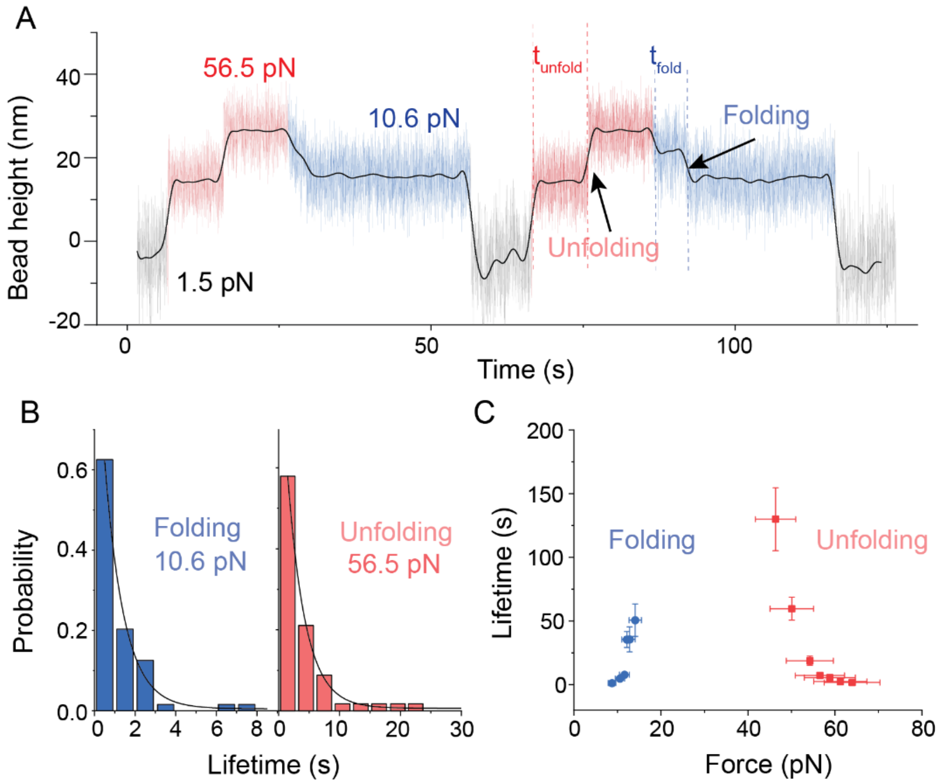


**Supplementary Figure 6. The mechanical responses of PF α-synuclein in force jumping experiments.** (A) A representative bead height−time trace containing unfolding and refolding events of PF α-synuclein in force-jumping cycles. The external force is jumping automatically from a resting force of 1.5 ± 0.15 pN to an unfolding force of 56.5 ± 5.7 pN, which is to allow the unfolding of the PF intermediate, and then jumping to a refolding force of 10.6 ± 1.1 pN, which is to allow the refolding of the PF intermediate. By repeating such force-jumping operations on multiple molecules for multiple cycles, the force-dependent unfolding and refolding lifetime of the PF intermediate at each force can be obtained. Raw data are indicated by gray (resting), red (unfolding), and blue (refolding), respectively. The smoothed data using FFT with 50 points of window is indicated by black. (B) Histograms of the unfolding lifetime (left panel) and refolding lifetime (right panel) at 56.5 ± 5.7 pN (n=57) and 10.6 ± 1.1 pN (n=59), respectively. An average unfolding lifetime of $\tau$ = 2.93 ± 0.11 s at 60.0 ± 6.0 pN and an average refolding lifetime of $\tau$ = 0.97 ± 0.09 s at 9.5 ± 0.95 pN were obtained by single-exponential function fitting (black curve), respectively. ± indicates S.E.. (C) Force-dependent average lifetime of unfolding (red) and refolding (blue) of PF α-synuclein. The average unfolding forces are at 46.4 ± 4.6 pN (n=6), 50.0 ± 5.0 pN (n=40), 54.2 ± 5.4 pN (n=13), 56.5 ± 5.7 pN (n=57), 60.0 ± 6.0 pN (n=57), 61.2 ± 6.1 pN (n=41), and 64.0 ± 6.4 pN (n=53), respectively. The average refolding forces are at 8.8 ± 0.88 pN (n=64), 10.6 ± 1.1 pN (n=59), 11.6 ± 1.2 pN (n=35), 12.2 ± 1.2 pN (n=28), 12.8 ± 1.3 pN (n=10), and 14.1 ± 1.4 pN (n=15), respectively. Error bars indicate the mean ± S.E..

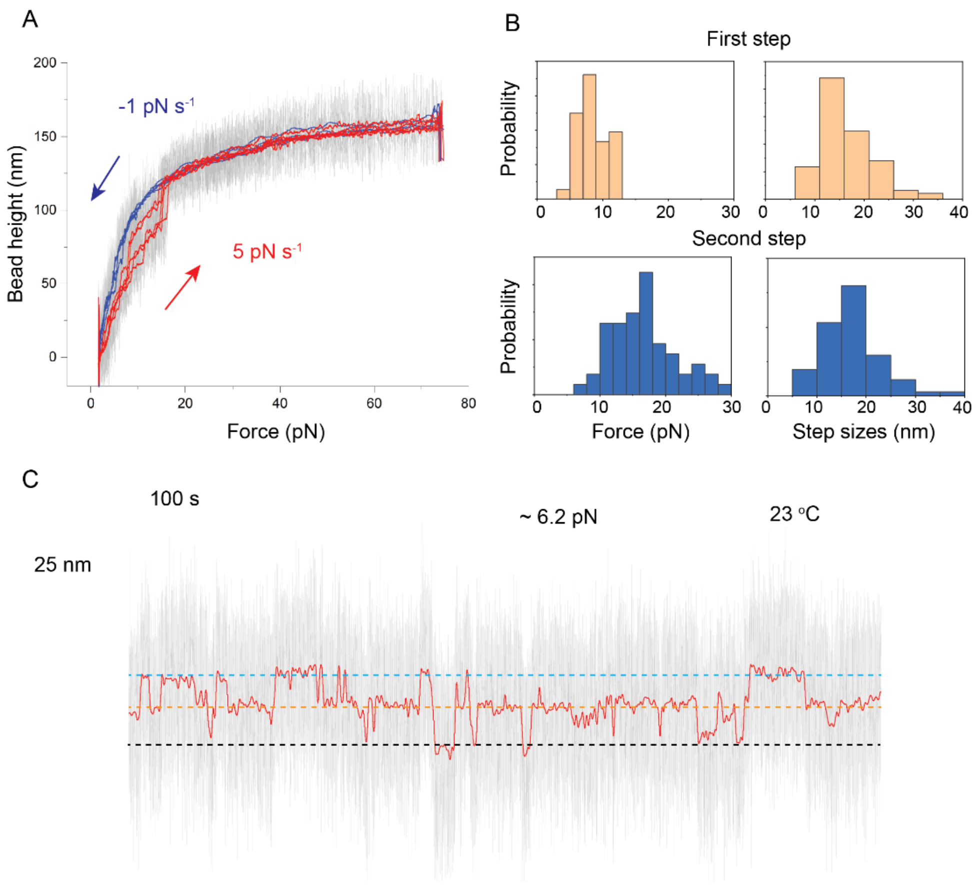


**Supplementary Figure 7. The mechanical responses of the minor species of α-synuclein.** (A) Representative force-extension curves in force loading rate scan showing a two-step unfolding of the minor species α-synuclein, where the stretching force is increased at a constant loading rate of 5 pN $s^{-1}$ and subsequently decreased to around 1.5 pN at a constant loading rate of −1 pN $s^{-1}$. Raw data, smoothed stretching curves, and smoothed relaxation curves (200 FFT) are presented as gray, red, and blue lines, respectively. (B) Unfolding force distributions and step size distributions of the two-step unfolding of the minor species α-synuclein at a loading rate of 5 pN $s^{-1}$. The numbers of observed first and second unfolding events are 36 and 54, respectively. (C) Representative bead height time trace at a constant force of 6.2 ± 0.6 pN shows a two-step dynamic fluctuation of the minor species α-synuclein. Raw data and 200-FFT-smoothed data are indicated by gray and red, respectively.

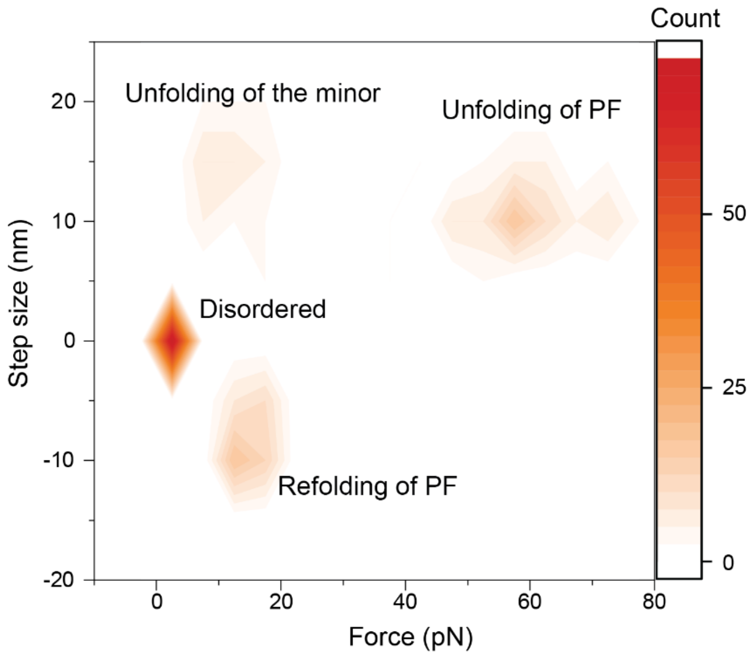


**Supplementary Figure 8. 2D map summarizes the mechanical responses of monomeric α-synuclein.** Molecule conformation fraction of N=131 monomeric α-synuclein molecules are based on their distinct unfolding and refolding signatures. The unfolding and refolding signatures of an α-synuclein were summarized based on their responses in the first ten force-scan cycles. Each count refers to a randomly chosen α-synuclein.

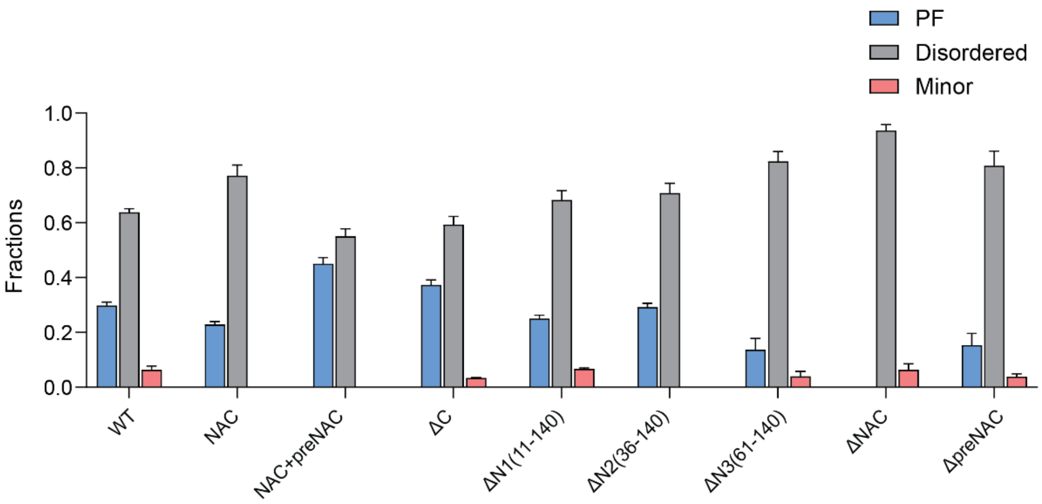


**Supplementary Figure 9. Species fractions of wild-type $\alpha$-synuclein and its truncations based on the unfolding signatures.** The total tested molecule numbers are: 131, 35, 20, 74, 60, 48, 51, 47, and 52 for WT, NAC, NAC+preNAC, ΔC, ΔN1, ΔN2, ΔN3, ΔNAC, and ΔpreNAC, respectively. Error bars indicate mean ± S.E.M.

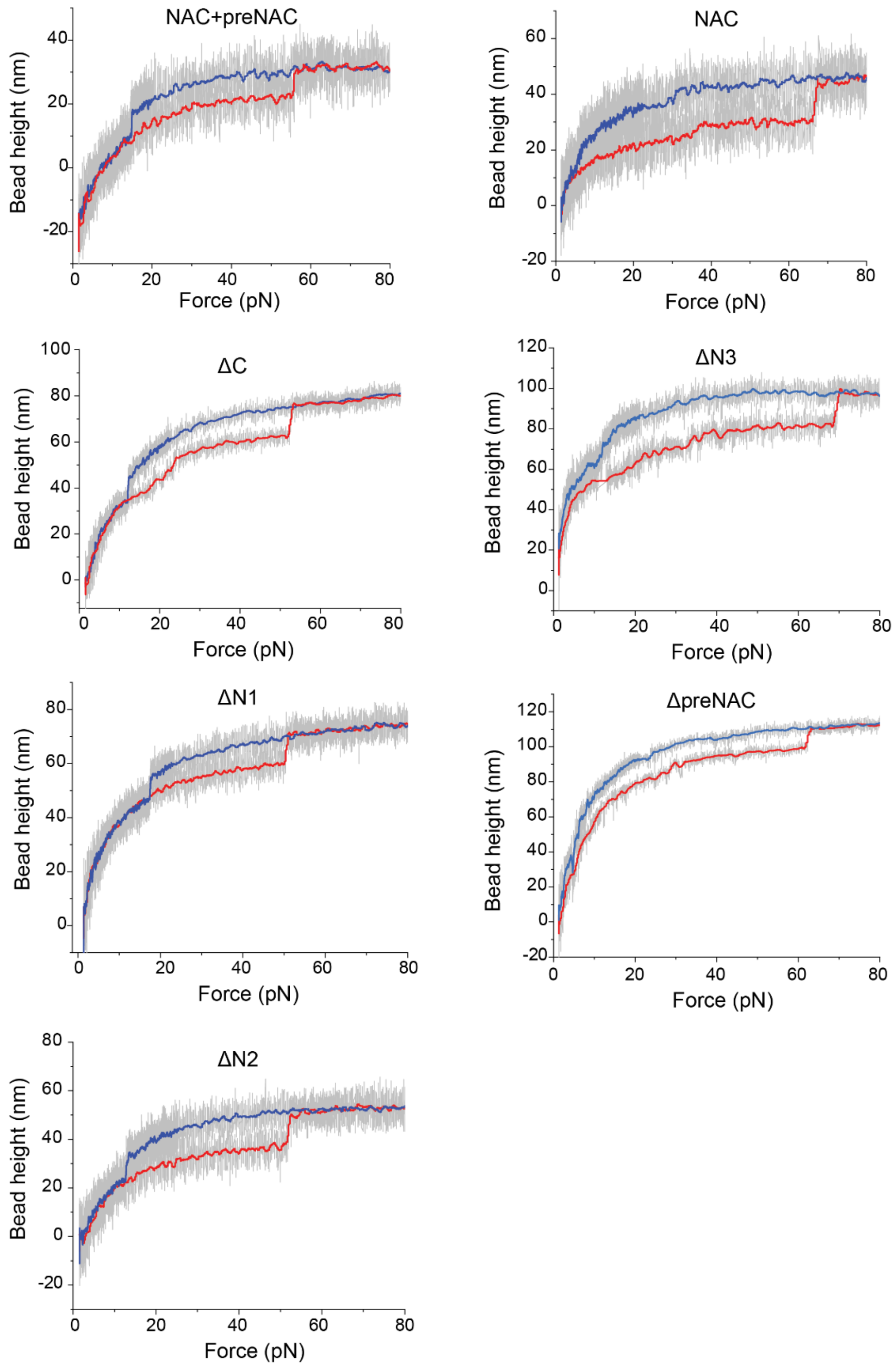


**Supplementary Figure 10. Typical force–bead height traces of $\alpha$-synuclein truncations with PF mechanical signature.** Raw data are indicated by grey. Smoothed stretching (5 pN $s^{-1}$) and relaxation (−1 pN $s^{-1}$) curves are indicated by red and blue, respectively.

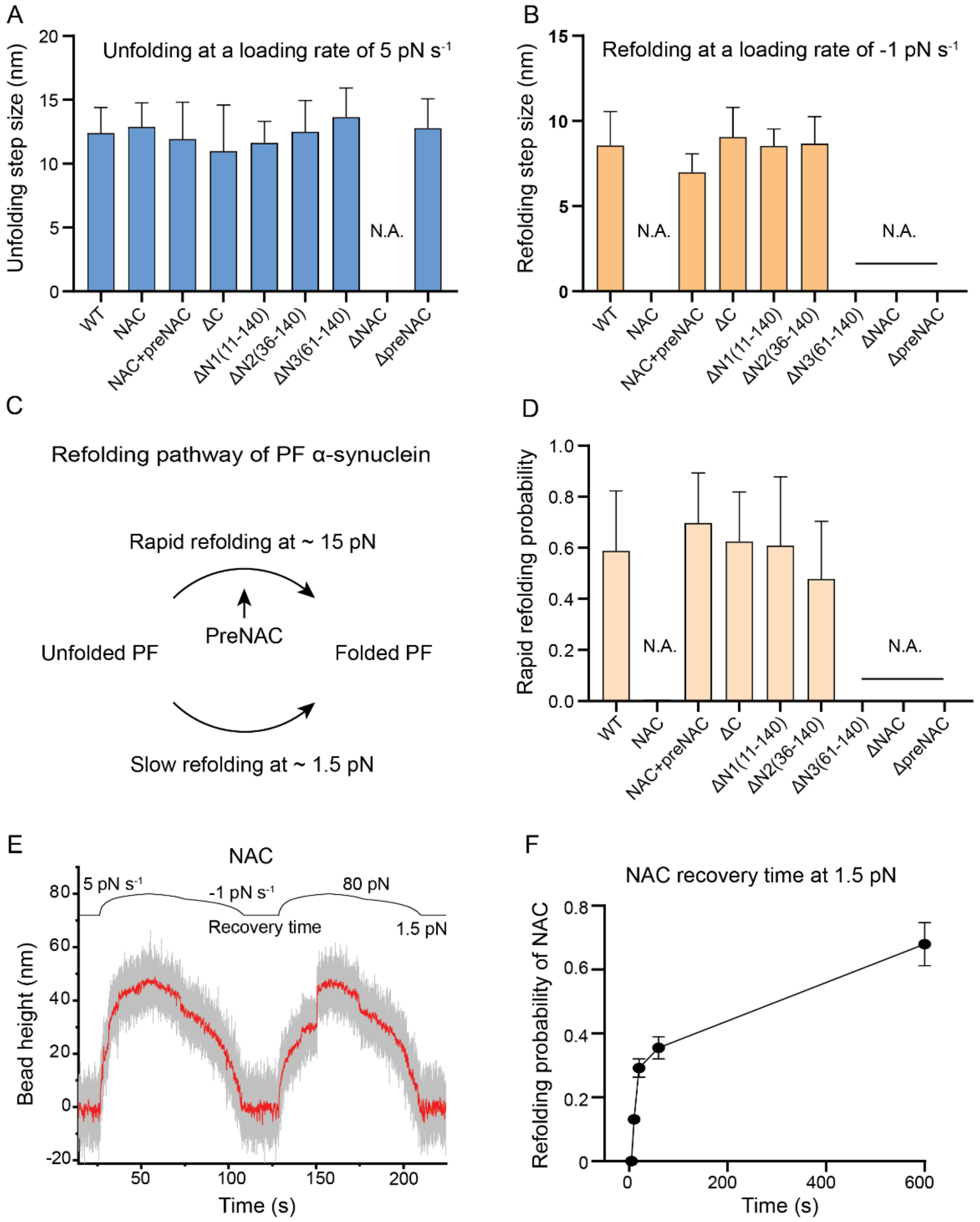


**Supplementary Figure 11. Mechanical signatures of truncated α-synuclein reveal that refolding of the unfolded PF is regulated by preNAC region.** (A) Unfolding step sizes of wild-type and truncated $\alpha$-synuclein proteins at a force-increasing loading rate of 5 pN s$^{-1}$. The number of unfolding events are: 124, 153, 193, 199, 161, 237, 16, 0, and 21 for WT, NAC, NAC+preNAC, ΔC, ΔN1, ΔN2, ΔN3, ΔNAC, and ΔpreNAC, respectively. Error bars indicate mean ± S.D.. (B) Refolding step sizes of wild-type and truncated $\alpha$-synuclein proteins at a force-increasing loading rate of -1 pN s$^{-1}$. The number of folding events are: 268, 0, 76, 55, 98, 89, 0, 0, and 0 for WT, NAC, NAC+preNAC, ΔC, ΔN1, ΔN2, ΔN3, ΔNAC, and ΔpreNAC, respectively. Error bars indicate mean ± S.D.. (C) Proposed refolding pathways of the unfolded PF. The rapid refolding observed at ~ 15 pN during the force decreasing from a high force at a loading rate of −1 pN s$^{-1}$, which is dominated by the existence of preNAC region. The slow refolding at ~ 1.5 pN is not dependent on the preNAC region. (D) Rapid refolding probability of the wild-type and truncated $\alpha$-synuclein proteins at a force-decreasing loading rate of −1 pN s$^{-1}$, which is quantified by the appearance of the folding stepwise bead height change at ~ 15 pN. N.A. means no such rapid refolding event was observed in all the experiments. (E) A representative time trace containing two force loading scanning cycles of NAC (61-100 a.a.) protein construct. Raw data and smoothed data using are indicated by gray and red, respectively. The external force is increased from ~1.5 pN to ~ 80 pN at a constant loading rate of 5 pN $s^{-1}$. Once the PF is unfolded, the force is decreased to ~ 10 pN at a constant loading rate of -5 pN $s^{-1}$ and subsequently decreased to ~ 1.5 pN at a slower force loading rate of -1 pN $s^{-1}$. The protein is then recovered at a resting force of ~ 1.5 pN for different waiting time durations. By repeating such force-scanning operations on multiple molecules for multiple cycles, the probability of the refolding (indicated by the bead height of the next round) after a waiting time at the resting force can be obtained. (F) Refolding probabilities of NAC were measured after waiting times of 0 s, 10 s, 30 s, 60 s, and 600 s at 1.5 pN, respectively, showing slow refolding of the NAC domain alone at ~1.5 pN. Error bars indicate mean ± S. E..

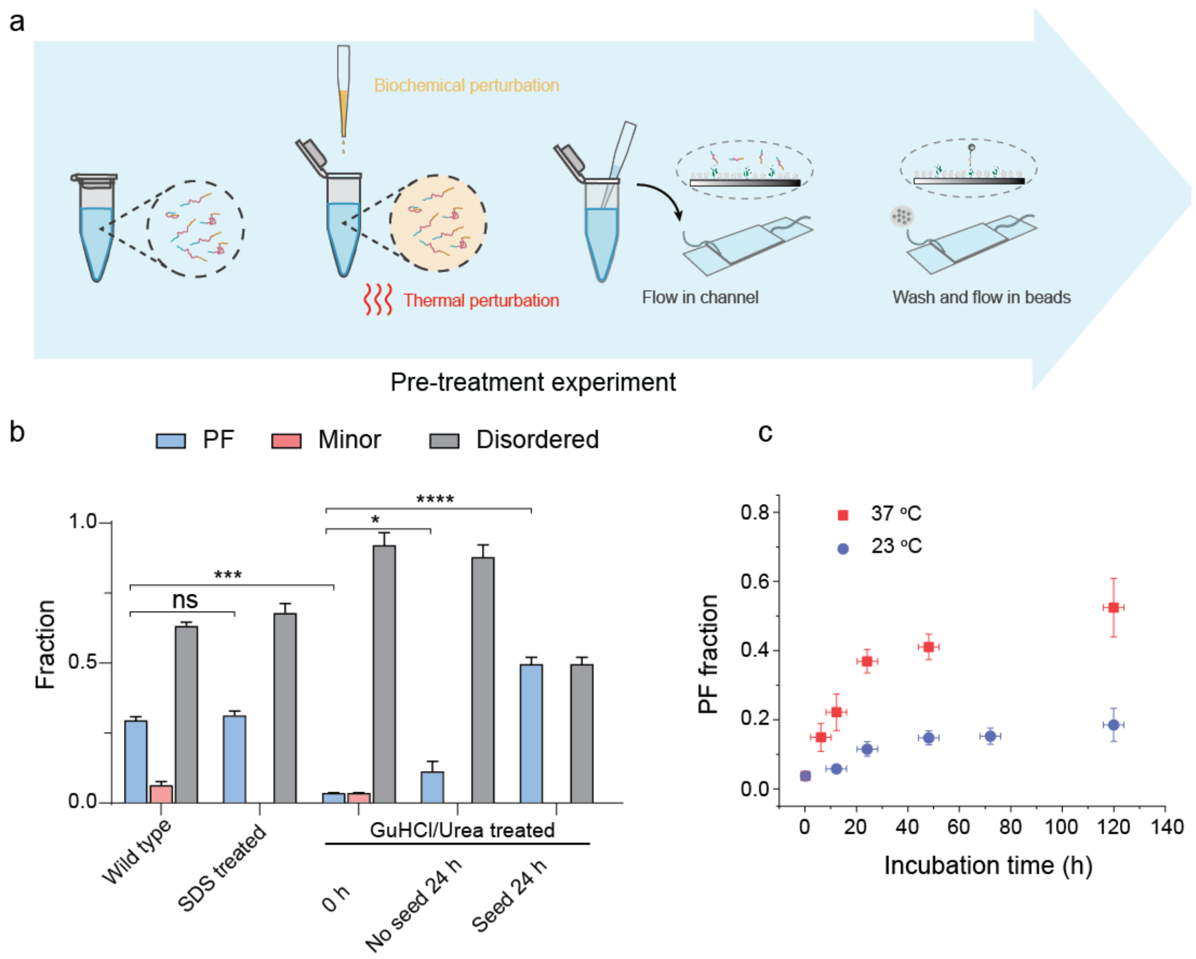


**Supplementary Figure 12. Effects of biochemical and thermal pre-treatments on the monomeric α-synuclein conformations.** (a) Schematic diagram of pre-treatment experiments, where the α-synuclein was subject to various biochemical reagents (denaturants, prion-like seed, chaperone, etc) or incubated at different temperatures before single-molecule analysis of the monomeric α-synuclein conformations. (b) Fractions of the three conformational species of α-synuclein (PF, disordered and minor conformations) after various pre-treatments: Wild type (untreated wild-type α-synuclein, N=131); SDS treated (2% SDS for 1 hour then removed, N=19); GuHCl/Urea treated (6 M GuHCl and 8 M urea for 1 hour then removed), followed by incubation at 23 °C for 0 hour (N=28) and 24 hours (N=44), or introduction with 1 μM prion-like seeds (small α-synuclein aggregates) at 23 °C for 24 hours (N=16). N is the number of the tested independent tethered molecules. Error bars indicate mean ± S.E.M.. (c) The PF fraction of GuHCl/urea pre-denatured α-synuclein after incubation of various time intervals up to 120 hours at 23 °C or 37 °C, respectively. Error bars indicate mean ± S.E.M.. ns: $p > 0.05$; *: $p < 0.05$; ***: $p < 0.001$; ****: $p < 0.0001$ (95% of confidence intervals). p values were determined by unpaired two-tailed Student's t-test.

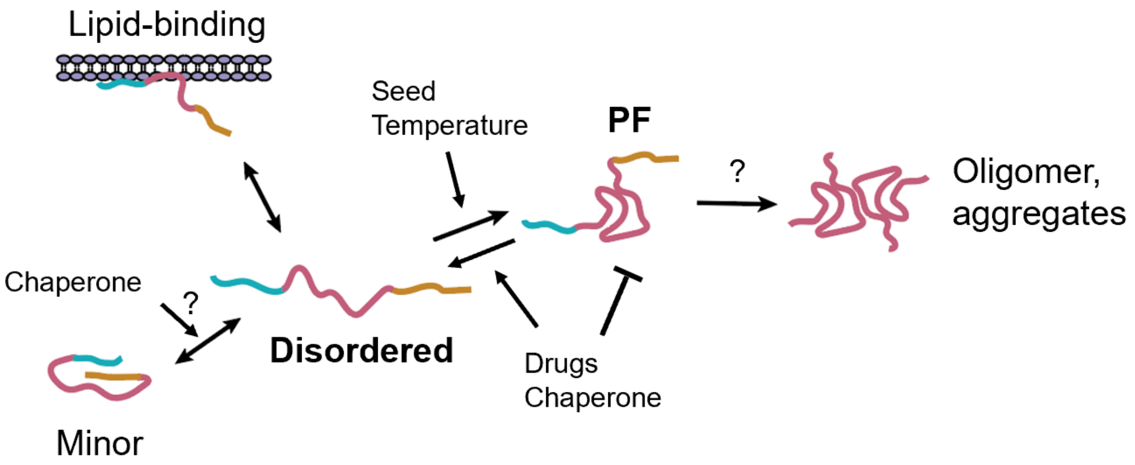


**Supplementary Figure 13. Illustrated diagram of the conformational structures and their transitions of monomeric α-synuclein.**
